# Age-dependent ventilator-induced lung injury: Mathematical modeling, experimental data, and statistical analysis

**DOI:** 10.1101/2023.04.20.537614

**Authors:** Quintessa Hay, Christopher Grubb, Sarah Minucci, Michael S. Valentine, Jennifer Van Mullekom, Rebecca L. Heise, Angela M. Reynolds

## Abstract

A variety of pulmonary insults can result in the necessity for mechanical ventilation, which, when misused, used for prolonged periods of time, or associated with an excessive inflammatory response, can result in ventilator-induced lung injury. Older patients have been observed to have an increased risk for respiratory distress with ventilation and more recent studies suggest that this could be linked to disparities in the inflammatory response. To address this, we ventilated young (2-3 months) and old (20-25 months) mice for 2 hours using high pressure mechanical ventilation and extracted data for inflammatory cell ratios, namely macrophage phenotypes, and lung tissue integrity. A large difference in naive macrophages at baseline, alternatively-activated (M2) macrophages at baseline, and airspace enlargement after ventilation was observed in the old mice. The experimental data was used to fit a mathematical model for the inflammatory response to lung injury. Model variables include inflammatory markers and cells, namely neutrophils and macrophages, epithelial cells at varying states, and repair mediators. Parameter sampling was performed using an iterative sampling method and parameter sets were selected based on their ability to fit either the old or young macrophage phenotype percentages and epithelial variables at zero and two hours. Classification methods were performed to identify influential parameters separating the old and young parameter sets as well as user-defined health states. Parameters involved in repair and damage to epithelial cells and parameters regulating the pro-inflammatory response were shown to be important. Local sensitivity analysis preformed for the different epithelial cell variables produced similar results. A pseudo-intervention was also performed on the parameter sets. The results were most influential for the old parameter sets, specifically those with poorer lung health. These results indicate potential targets for therapeutic interventions prior to and during ventilation, particularly for old subjects.

**Author summary:** A variety of inhaled pathogens and other pulmonary insults prompt the need for mechanical ventilation; a procedure that has become increasingly necessary following the 2019 coronavirus pandemic. A proportion of patients respond poorly to ventilation, some resulting in ventilator-induced lung injury. Observational data has shown increased instance of severe disease in older patients as well as differences in the inflammatory response to injury, although more research is needed to confirm this. We performed high-pressure ventilation on young (2-3 months) and old (20-25 months) mice and observed large disparities in inflammatory cell ratios at baseline and lung tissue integrity after ventilation. The experimental data was then used to fit a mathematical model of the inflammatory response to lung injury. We used a variety of analysis methods to identify important parameters separating the young and old parameter sets and user-defined health states of the resulting simulations. Parameters involved in damage and repair of epithelial cells in the lung as well as parameters controlling the pro-inflammatory response to injury were important in both classifying between old and young sets and determining predicted health after ventilation. These results indicate potential targets for therapeutic interventions prior to and during ventilation.

## Introduction

A variety of inhaled pathogens and other pulmonary insults illicit an immune response that causes inflammation in the lung tissues. Intense or persistent inflammation can damage the delicate alveolar tissue and result in acute respiratory distress syndrome (ARDS). This can progress to complete respiratory failure in some cases. To increase the probability of survival, the clinical intervention for ARDS is the use of mechanical ventilation (MV) [1]. While MV is often a necessary procedure, prolonged use or misuse of the ventilator may result in ventilator-induced lung injury (VILI). The damage caused to alveolar sacs (clusters of alveolar cells) during MV can lead to volutrauma (extreme stress/strain), barotrauma (air leaks), atelectrauma (repeated opening and closing of alveoli), and biotrauma (general severe inflammatory response). The culmination of these injuries can result in ventilator dependence, multi-system organ failure, or even death [2, 3].

Although VILI can occur in patients regardless of lung health [2], there is an observable disparity in the inflammatory response of older individuals as well as a higher incidence of critical disease [4–6]. Past research has reported increased levels of circulating inflammatory cytokines and altered macrophage function in older mice [7]. *In vivo* experimental results have also indicated increased risk of lung injury following ventilation for older mice [8]. These results were observed again in Herbert *et al.* [9]. Most recently, infections associated with the novel coronavirus have also exhibited an increased risk of mortality and severe disease in older patients [10]. One particular study found that 6.6% of participants aged 60 years of age and older developed critical disease following a SARS-CoV-2 infection; this is approximately twelve times higher than in younger participants (0.54%) [6]. These observed discrepancies in the inflammatory response and increased rate of mortality and severe disease in elderly patients stress the need for further studies of VILI in regards to aging.

Nin *et. al* [11] found increased susceptibility and severity to pulmonary and vascular dysfunction associated with age during high tidal volume ventilation in mice. Older mice also exhibited increased levels of inflammation marked by a higher concentration of interleukin-6, a pro-inflammatory cytokine, and aspartate aminotransferase, a non-specific marker of cell injury. This intrinsic decline in the effectiveness of the innate immune response has been studied extensively [12–15]. Most notably, Dace and Apte [15] found that aging affected the polarization of macrophages, an immune cell that can exhibit a range of pro-and anti-inflammatory properties. An effective immune response relies on both a pro-inflammatory response to rid the insult of foreign cells and other debris and an anti-inflammatory response to regulate the pro-inflammatory response, promote repair, and remove debris incurred by early-response phagocytes. With aging, polarization of macrophages was observed to be skewed toward the alternatively-activated, M2, phenotype. Decreased activation of the classically-activated, M1, phenotype, generally associated with a pro-inflammatory response, could result in a decreased ability to clear infections, thus prolonging the inflammatory response and inhibiting later stages of healing.

An imbalance in the pro- and anti-inflammatory responses can cause additional complications for the individual during various injuries and insults. Macrophages in particular play a significant role in the impact of aging on the immune response [7, 16, 17]. Therefore, to develop interventions to mitigate the effects of VILI, it is important to study the immune response to lung injury and the interplay between various types of cells. We are focused on the innate immune cells, neutrophils and macrophages, their associated cytokines, and the alveolar epithelium, which consists of alveolar type I and type II cells. Alveolar type I cells make up about 95% of the alveolar surface and are primarily responsible for facilitating gas exchange. Type II cells cover the other 5% of the surface and are important in the innate immune response. In the presence of damage, these cells proliferate to repair the epithelium and can also differentiate to type I cells [18, 19]. In the present study, we examine these cells in 2-3 month old mice (young) and 20-25 month old mice (old) exposed to high-pressure MV for up to 2 hours. We present broad macrophage sub-phenotypes, M1 and M2, obtained from flow cytometry and quantitative measures of lung damage at the alveolar epithelial-endothelial barrier.

We use mathematical modeling and statistical methods to investigate the differences in the pulmonary innate immune response and predicted outcome for young and old versions of the model. At this stage of exploration of VILI, we focus on epithelial damage and immune system interactions in young and old mice. It is difficult to clinically isolate the local epithelial and inflammatory response in the lung during VILI and often expensive to collect quality data. For this reason, we rely on *in silico* modeling of experimental data to supplement the available *in vivo* data. These *in silico* approaches provide insight into the immune response and the nonlinear dynamics of the system. The resulting analysis is used to identify important factors and generate hypotheses [20].

### Mathematical Background

The immune response to bacterial and viral infections has been studied extensively [21–26]. More recent models have focused specifically on COVID-19 infections [27–31]. Models for general lung injury not caused by infection have also been explored [32–35] as well as general inflammatory dynamics [4, 36]. A review of mathematical models that focus generally on the immune response in the lungs has also been published [37]. Past models for MV and VILI have generally focused on airway mechanics to inform and optimize machine settings and assess stress [38–46], with some models incorporating fluid-structure interactions [44, 47, 48]. Models have also been developed to include epithelium and immune cells [47, 49] and modified to include a response to an epithelial wound [50], an infection [51, 52], and other applications. [53, 54].

Torres *et al.* modeled the innate immune response to bacteria and macrophage polarization along the pro-to anti-inflammatory spectrum [55]. Minucci *et al.* included these interactions as well as epithelial dynamics and damage-induced recruitment of immune cells. An epithelial subsystem was combined with the Torres *et al.* model and ventilator-induced epithelial cell damage triggered the immune response rather than an infection. The current model is an expansion of the model built in Minucci *et al.* by including terms modeling epithelial barrier breakdown leading to increased cytokines and immune cells in the alveolar compartment [56]. The resulting model has 19 variables and 64 parameters. In this study we coupled this model with the young and old experimental data to explore age-dependent outcomes and dynamics.

These mechanistic, equation-based models are often used in conjunction with statistically-based methods and models [57] to understand the possible dynamics associated with varying parameters sets. Parameter sampling and post-analysis of the mechanistic data obtained from the model are just two examples of processes that benefit from a statistical approach. To sample large parameter spaces, numerous aptly named ‘space-filling designs’ have been developed since the advent of computer experiments in the 1970s. Perhaps the most commonly used design is Latin hypercube sampling [58], but many others are used, including uniform sampling and maximin designs [59]. Many others have built off these general designs that work only for continuous data, and created variants and alternatives more specifically geared towards individual use-cases, such as the sliced latin hypercube design [60] for categorical inputs and the fast flexible filling algorithm [61], designed for non-rectangular spaces. Machine learning algorithms have also aided in the analysis of mechanistic models. Methods such as random forest, neutral networks, and principal components analysis continue to be used in congruence with mathematical models and biological systems [62–66]. These methods work well to process the large amounts of data obtained from a mechanistic model and identify abstract features of the system [57]. These algorithms can also identify nonlinear interactions between factors within the model, adding crucial insight into parameters effecting the response.

Sensitivity analysis is a useful tool for models with a large number of parameters where baseline values are unknown or difficult to measure. These methods analytically measure changes to the model outputs from perturbations in the model inputs [67] and include local and global methods. In local sensitivity analysis, the model output is observed when only one model parameter is varied around a selected nominal value and all other parameters are held constant. Global methods examine the sensitivity of parameters within the entire parameter space. Global techniques are usually implemented using Monte Carlo simulations, giving them the description of sampling-based methods [68]. This includes methods like Pearson correlation coefficient and partial correlation coefficients for linear trends and Spearman rank correlation coefficient and partial rank correlation coefficient for nonlinear trends with a monotonic relationship between inputs and outputs. Nonlinear non-monotonic trends require slightly different methods based on decomposition of model output variance. These methods include the Sobol method and its extended version based on (quasi) random numbers and an *ad hoc* design [69], and the Fourier amplitude sensitivity test and its extended version. The methods have been implemented in various models involving wound healing and the inflammatory response [70–74]

To determine plausible parameter sets for our model using both the young and old experimental data, we initially sampled using a beta distribution to favor lower parameter values, and then performed an iterative stochastic local search around likely candidates. The parameter sets were then simulated and equation values were compared with *in vivo* data to determine old or young presenting behavior in the resulting transients before and after ventilation. Various transient features were also calculated and analyzed including epithelial qualities and inflammatory cell quantities. Analysis of the resulting parameter sets included identifying parameters associated with old or young data, analyzing differences in lung health states between the old and young sets, and determining parameters associated with poorer lung health both including and excluding age classification. Further investigation of the identified parameter sets included local sensitivity analysis to assess model output sensitivity to variations in the parameters. The results from classification and sensitivity analysis were used to simulate pseudo-interventions to determine parameters that may be modulated to improve epithelial health during MV. This can help inform potential therapeutic targets for patients that are considered high risk before ventilation or even patients that present signs of distress during ventilation.

## Materials and methods

### Ethics statement

Animal Research (involving vertebrate animals, embryos or tissues) Male C57BL/6 mice 8 weeks of age were purchased from Jackson Laboratory (Bar Harbor, ME). All animals were housed in accordance with guidelines from the American Association for Laboratory Animal Care and Research protocols and approved by the Institutional Animal Care Use Committee at Virginia Commonwealth University (Protocol No. AD10000465).

### Experimental materials & methods

#### Animals

Male C57BL/6 mice 8 weeks of age were purchased from Jackson Laboratory (Bar Harbor, ME). Male C57BL/6 mice 20 months of age were provided by the National Institute on Aging (Bethesda, MD). All animals were housed in accordance with guidelines from the American Association for Laboratory Animal Care and Research protocols and approved by the Institutional Animal Care Use Committee at Virginia Commonwealth University (Protocol No. AD10000465).

#### Pressure-controlled ventilator-induced lung injury model

We mechanically ventilated young (2-3 months) and old (20-25 months) C57BL/6J wild-type mice using a Scireq FlexiVent computer-driven small-animal ventilator (Montreal, Canada) and previously cited methods in Herbert *et al.* [9] with slight modifications. Mice were anesthetized, tracheotomized, and then ventilated for 5 minutes using a low pressure-controlled strategy (peak inspiratory pressure (PIP): 15 cmH20, respiratory rate (RR): 125 breaths/min, positive end-expiratory pressure (PEEP): 3 cmH20). Mice were then ventilated for 2 hours using a high pressure-controlled mechanical ventilation (PCMV) protocol (PIP: 35-45 cmH20, RR: 90 breaths/min, and PEEP: 0 cmH20). Pulmonary function and tissue mechanics were measured and collected at baseline and every hour during the 2-hour high PCMV duration using the SCIREQ FlexiVent system and FlexiWare 7 Software. A separate group of mice was anesthetized, tracheotomized, and maintained on spontaneous ventilation for 2 hours.

### Tissue processing

Immediately following mechanical ventilation, the right lobes of the lung were snap frozen with liquid nitrogen, then stored at -80*^◦^*C for further analysis. The left lobes of the lung were then inflated with digestion solution containing 1.5 mg/mL of Collagenase A (Roche) and 0.4 mg/mL DNaseI (Roche) in HBSS with 5% fetal bovine serum and 10mM HEPES and processed as previously described Yu *et al.* [75]. The resulting cells were counted, and dead cells were excluded using trypan blue. Subsets of the experimental groups were also used to collect left lobes for histological analysis.

#### Histological analysis

Lung tissue samples were embedded and stained with hematoxylin and eosin (H&E). The mean linear intercept, an index of airspace enlargement, was used to quantify relative differences in alveolar airspace area within lung histology sections. These were measured and analyzed as previously described Herbert *et al.* [9].

#### Flow cytometric analysis

Following live cell counts, 4*×*10^6^ cells per sample were incubated in blocking solution containing 5% fetal bovine serum and 2% FcBlock (BD Biosciences) in PBS. The cells were then stained using a previously validated immunophenotyping panel of fluorochrome-conjugated antibodies [76] with slight modifications (See S1 Fig. for a list of antibodies, clones, manufacturers, and concentrations). Following the staining procedure, cells were washed and fixed with 1% paraformaldehyde in PBS. Data were acquired and analyzed with a BD LSRFortessa-X20 flow cytometer using BD FACSDiva software (BD Bioscience). Histogram plots were generated using FCS Express 5 software (De Novo). Compensation was performed on the BD LSRFortessa-X20 flow cytometer at the beginning of each experiment. “Fluorescence minus one” controls were used when necessary. Cell populations were identified using a sequential gating strategy that was previously developed in Misharin *et al.* [76]. The expression of activation markers was presented as median fluorescence intensity (MFI).

### Mathematical model and analysis methods

#### Model equations

The model uses differential equations to track the transition from a healthy lung state to a state with damage to the epithelial cells in response to ventilation. This a direct transition from a healthy state to damaged state at the cellular level, we do not explicitly model the stress and strains at the tissue level that give rise to this damage. In our model damaged cells produce mediators that activate innate immune cells, neutrophils and macrophages. Immune cell influx causes additional damage to the lung epithelial cells. The epithelial cells can 1) return to a health state via repair, which is regulated by repair mediators, 2) transition to the death/empty state. The portion of the population that is in the death/empty state represents the portion of the lung that needs to be replaced via proliferation of healthy epithelial cells. A full model schematic including the dynamics is given in Fig. 1. Model variables are in Table 1 and the parameters with brief descriptions are in Table 2.

**Fig 1.**
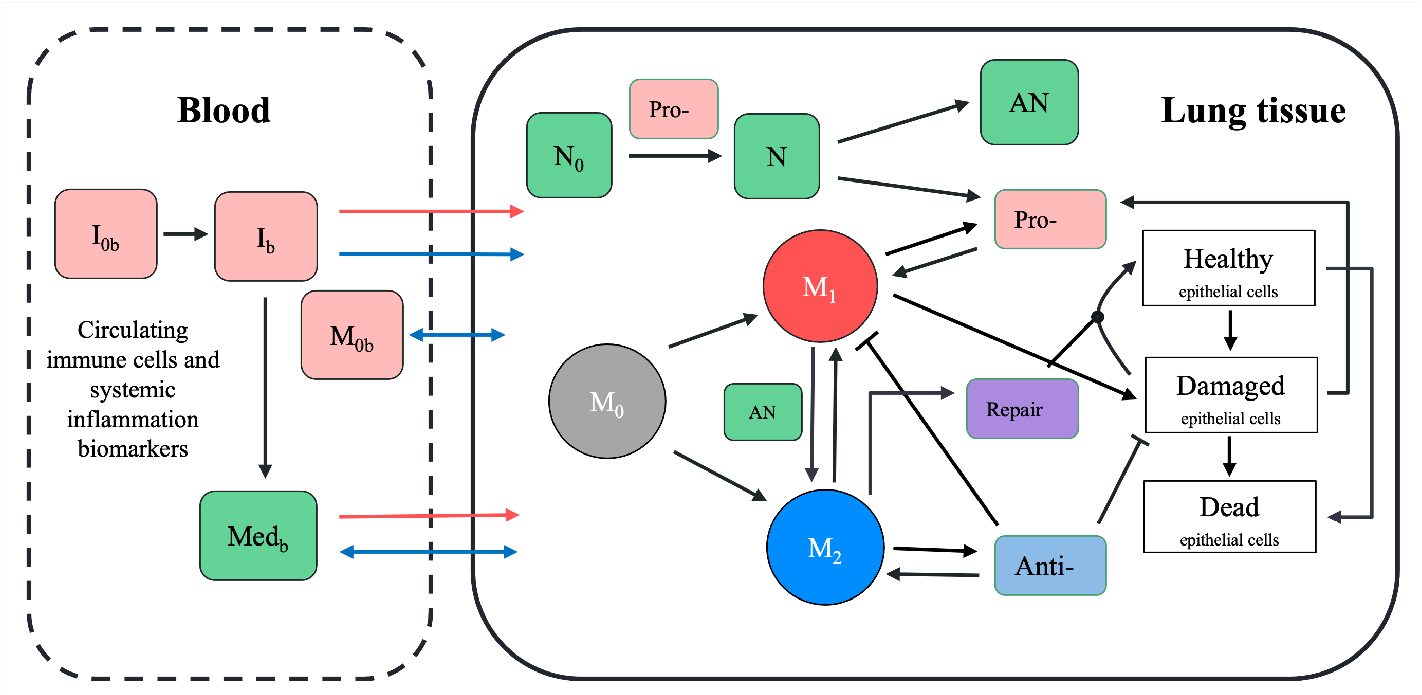
Model Schematic. The model has two compartments: lung tissue and blood. The various circles and boxes represent the different inflammatory cells, mediators, and epithelial cell states. The variables *I*_0*b*_ and *I_b_*in the blood compartment represent an unactivated or undifferentiated and an activated or differentiated general inflammatory cell, respectively. This includes the various states of neutrophils and macrophages. The variable Med*_b_* in the blood compartment represents both pro- and anti-inflammatory mediators in the blood. Black arrows represent upregulation or transition and black lines with a perpendicular edge represent inhibition or down-regulation. The blue arrows represent movement between the two compartments, either diffusion based or at a constant rate. The red arrows represent movement from the blood into the lung compartment as a result of epithelial barrier degradation.

**Table 1.** Table of model variables with descriptions.

| Bloodstream | Lung | Description |
| --- | --- | --- |
| | $E_h$ | Healthy epithelial cells |
| | $E_d$ | Damaged epithelial cells |
| | $E_e$ | Dead epithelial cells/empty space |
| $p_b$ | $p$ | Pro-inflammatory mediators |
| $a_b$ | $a$ | Anti-inflammatory mediators |
| $M_{0b}$ | $M_0$ | Naive macrophages |
| $M_{1b}$ | $M_1$ | M1, classically-activated macrophages |
| $M_{2b}$ | $M_2$ | M2 alternatively-activated macrophages |
| $N_{0b}$ | $N_0$ | Unactivated neutrophils |
| $N_b$ | $N$ | Activated neutrophils |
| | $AN$ | Apoptotic neutrophils |
| | $R$ | Repair mediators |

$E_e$ is fit to experimental data as a proportion of epithelial space in the lung tissue. The epithelial variables $E_d$ and $E_h$ are also proportions, such that $E_h + E_d + E_e = 1$ . The remaining variables have arbitrary units for simulation. The variables $M_0$ , $M_1$ , and $M_2$ are used to calculate percentages for each phenotype, which is then fit to the experimental flow cytometry data after simulation as proportional values.

**Table 2.**
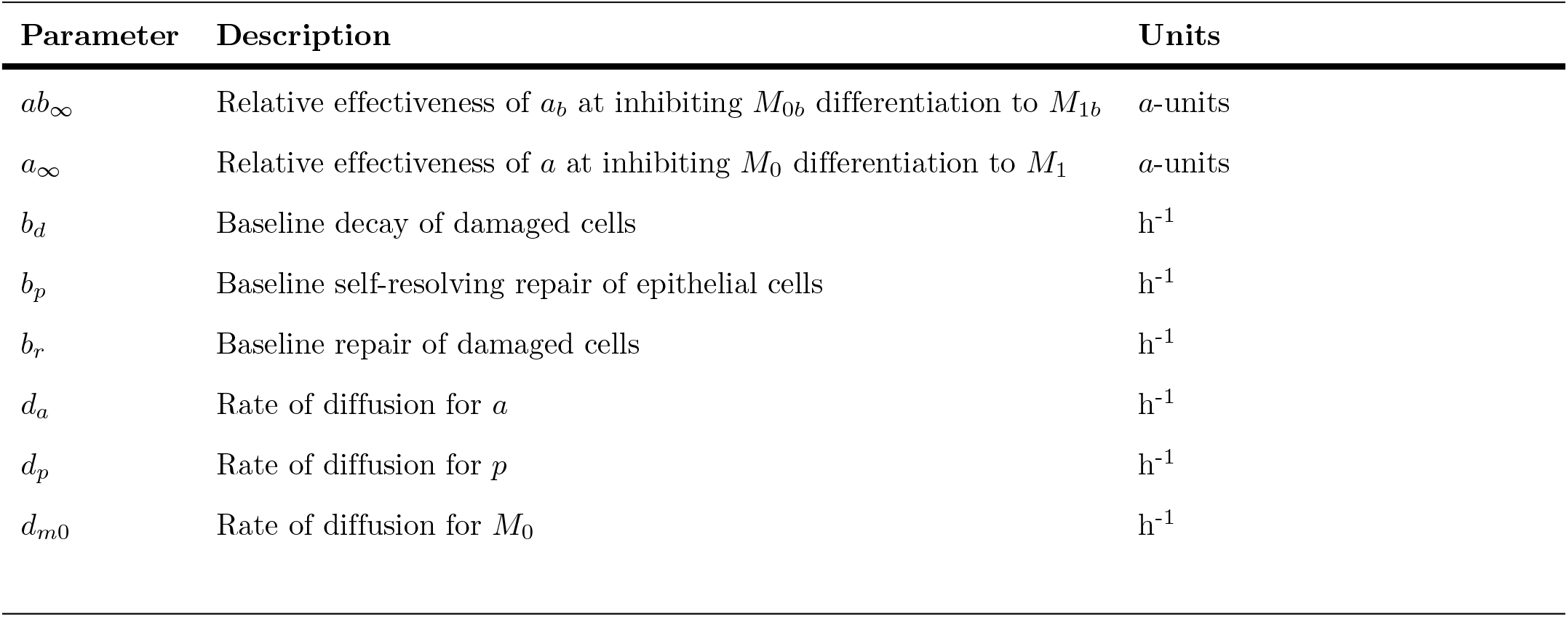
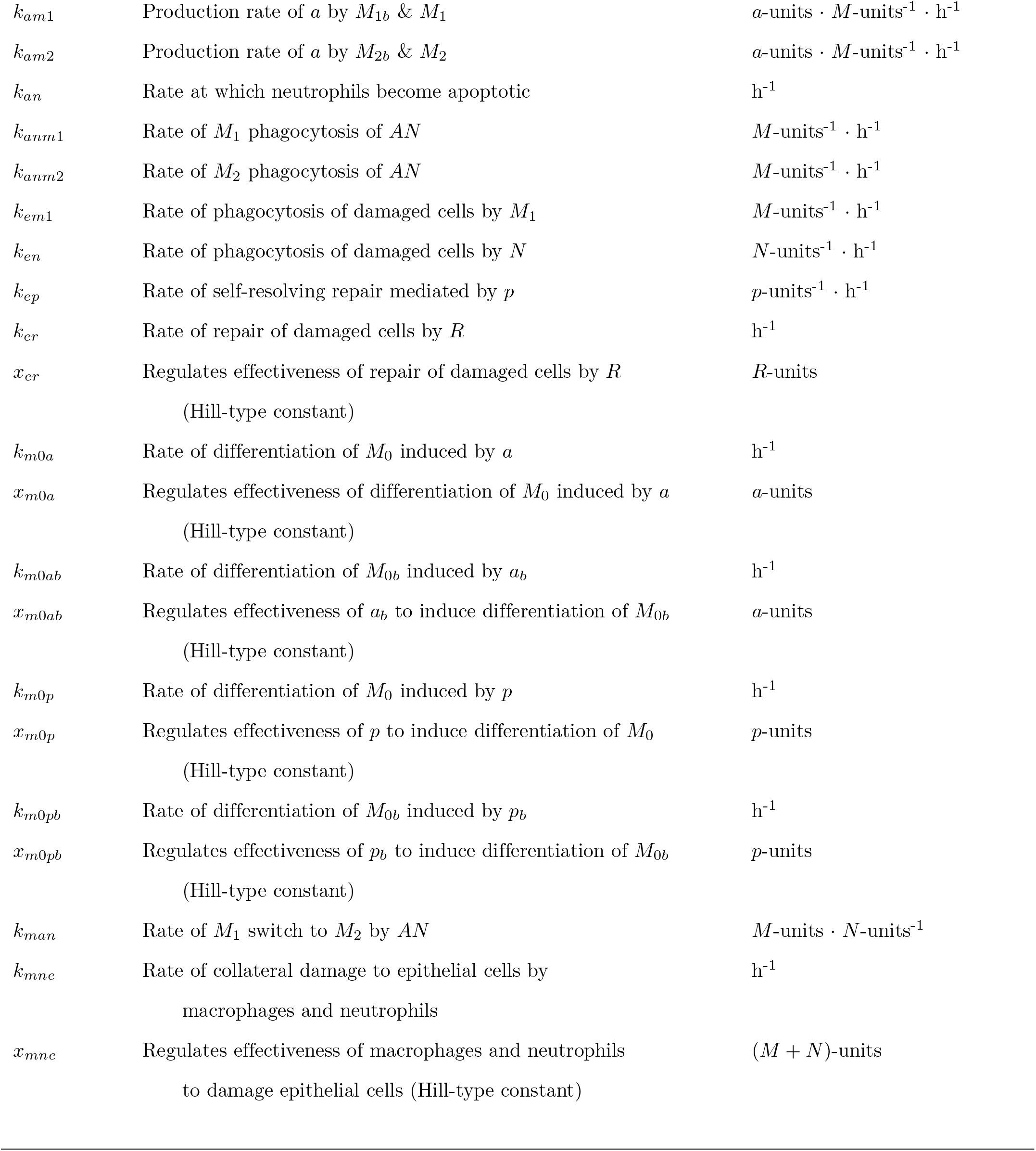
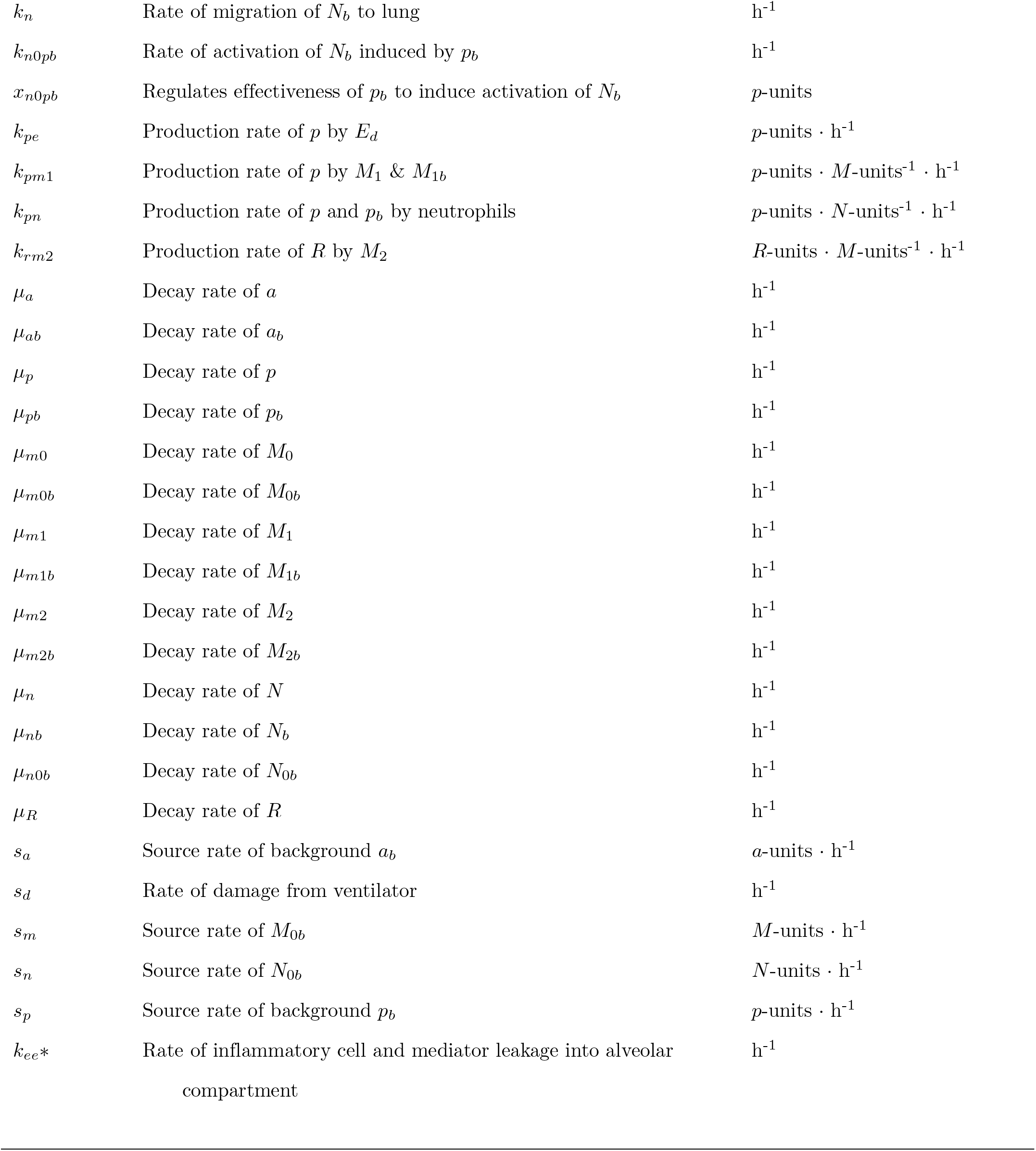
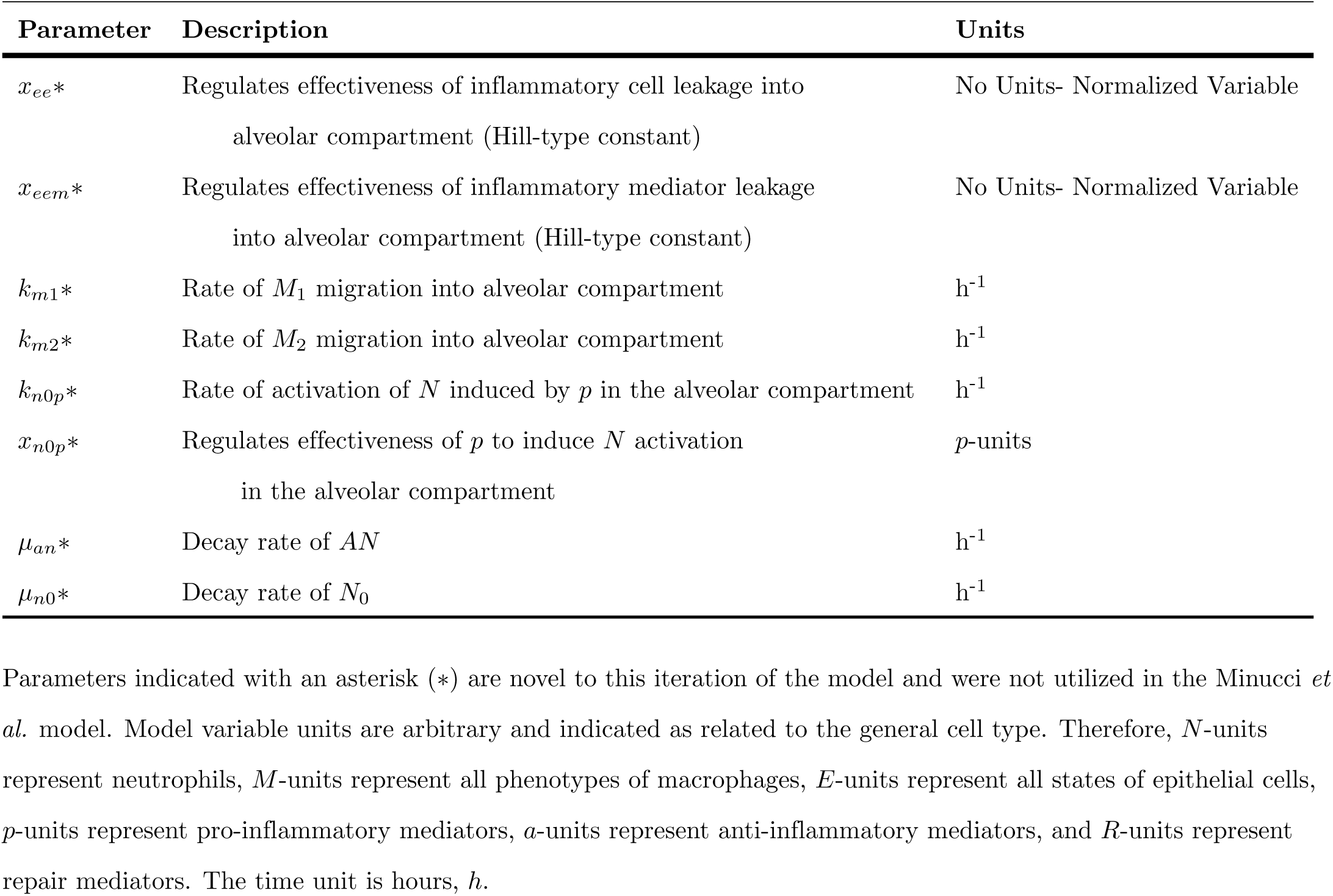
Model parameters with descriptions.

We account for macrophage phenotype on a population level. Therefore, our variables track the overall level of M1 type activity (classically activated) versus M2 type activity (alternatively activated). Activation of naive macrophages M0 by pro-inflammatory mediators (PIMs) give rise to the M1 phenotype, phagocyte cells producing PIMs and anti-inflammatory mediators (AIMs). M1 cells phagocytize both damaged epithelial cells and apoptotic neutrophils. Neutrophils also phagocytize damaged epithelial cells and produce PIMs.

Conversely, activation of naive cells by AIMs gives rise to a M2 phenotype, producing AIMs and repair mediators. M1 cells can transition to M2 cells in response to phagocytising apoptotic neutrophils. The full equations for our model are in Appendix S2 - S7. The main reference for the equation derivations are given in Minucci *et al.*, since our current model is an adaptation of the Minucci *et al.* model.

The main change from Minucci *et al.* in this model was the introduction of a breakdown in the barrier integrity that leads to movement between the systemic blood compartment into the lung space that is not diffusion driven. We illustrate this with an immune cell equation, the lung *M*_0_ equation (Eqn. 1), and a mediator equation, the lung PIMs (*p*, Eqn. 2), but this type of term occurs for all the immune cells and mediators (see Appendix S2 - S6). The second to last term in the *M*_0_ equation (Eqn. 1) is a nonlinear Hill-type term that allows naive cells to move from the blood into the lung when the epithelial lung barrier is degraded significantly. Therefore, the term is dependent on *E_e_* with the parameter *x_ee_* controlling the level of *E_e_*at which this term achieves its half max. *x_ee_* is fixed to 0.75 for all immune cell equations. The second to last term of the *p* equation (Eqn. 2) has the same form but with the parameter defined as *x_eem_*. *x_eem_* is fixed to 0.5 for all mediator equations. These parameters are fixed at these levels to ensure that the smaller mediators pass through the degraded alveolar-capillary barrier at lower *E_e_* level than the immune cells. This fixed level is also a first step to mapping the epithelial variables to clinically relevant lung injury, such as pulmonary edema. Mediator flux is associated with severe lung damage (such as edema) and immune cell leakage into the lung due to barrier disruption which would lead to ventilation failure as the alveoli fill with fluid instead of air. The third from last term in both of these equations model the diffusion driven transition between compartments.

The first term in the *M*_0_ equation (Eqn. 1) models the activation of the naive macrophages into the M1 and M2 phenotypes with down-regulation of the M1 phenotype via inhibition by AIMs represented by the variable *a*. The first three terms of the *p* equation (Eqn. 2) models production of PIMs by damaged epithelial cells, M1 cells (inhibited by *a*) and neutrophils, respectively. The last term in both equation models intrinsic decay.

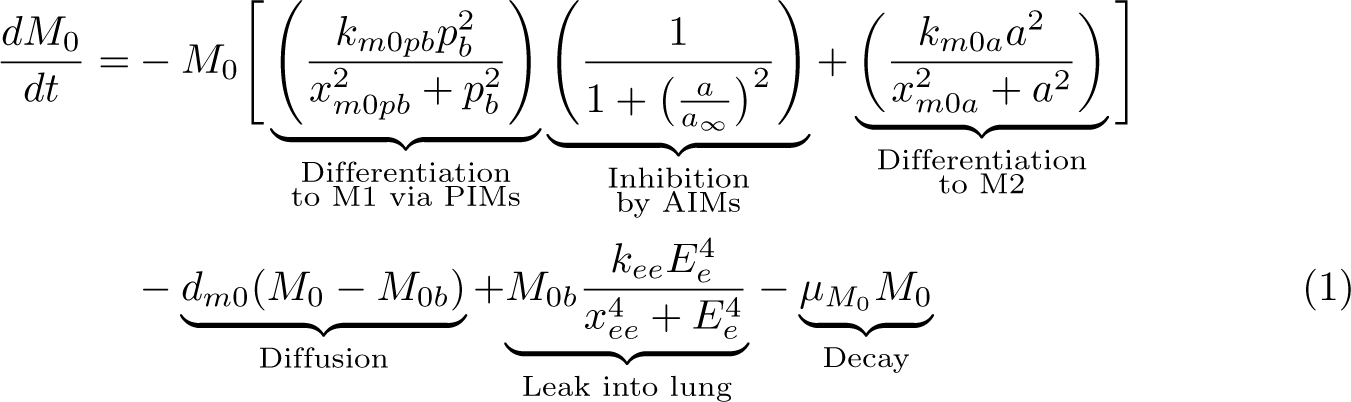

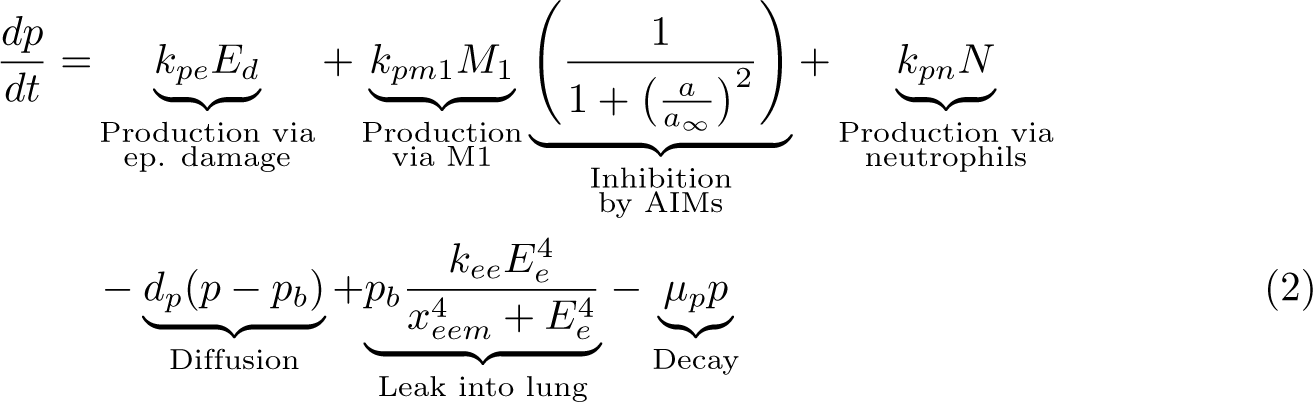

#### Sampling and parameter selection

The parameters sets were simulated using MATLAB (R2021a) [77] to determine plausible *in silico* transients associated with each experimental group. Plausible parameters sets are those that reached numerical steady state and fell with in the range of *in vivo* data for either the young or old data at each time point. Transients were compared with the experimental data for unactivated macrophages (*M*_0_), classically-activated macrophages (*M*_1_), alternatively-activated macrophages (*M*_2_), and airspace enlargement (*E_e_*).

The data ranges used for each experimental group and cell are plotted in Fig. 2. Note from Fig. 2 some data ranges overlap significantly, but given there is separation in the M0 and M2 data for time zero there are no parameter sets that satisfy both young and old conditions. Past experiments have observed differences in the inflammatory response in older individuals [12–15]. The experimental data collected exhibited these differences in the varying activation of macrophages. There was also an observable difference in the airspace enlargement variable after ventilation shown in Fig. 2. The observed ranges do overlap, but percentages tended to be larger in the old group.

**Fig 2.**
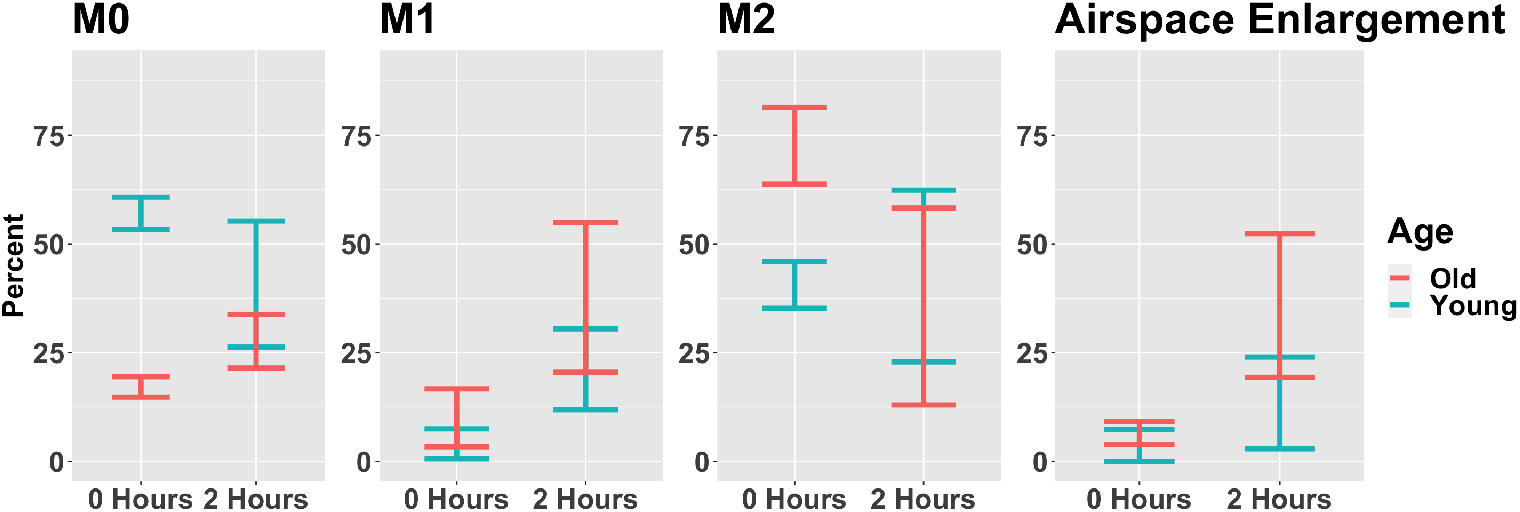
Summary of data ranges used for parameter fitting. The plot gives the range of experimental values used to assess the numerical simulations for biological plausibility. Each range consists of 3-6 experimental observations since some data points were excluded as outliers. Outliers were defined as those being more than two standard deviations outside the mean.

Parameter sets were first simulated in MATLAB [77] to reach numerical steady-state to ensure that the observed dynamics were only a product of the ventilation. Parameter sets were simulated with the parameter *s_d_*, a parameter used to describe the damage caused by ventilation, set to zero. Each parameter set was simulated with up to two possible initial conditions for 800 hours and classified as approaching numerical steady state if the Euclidean norm of the difference between the end values of the variables and the value at each numerical time step within the last 20 hours of simulation was less than 0.001. Parameter sets that achieved steady state and fit either the young or old initial data values were then simulated for 200 hours with ventilation, *s_d_ >* 0, for the first two hours. Parameters sets then compared against the data at 2 hours and classified as either young, old, or neither depending on the resulting transients.

Candidate parameter sets for this process were found using a three-step parameter sampling process in R [78]. In the initial step, a large number of samples were generated using a scaled *Beta*(1, 3) distribution with a large scale parameter to sample between 0 and 120 (to include multiple orders of magnitude). Parameter sets with associated transients that fell within the range of the data for all macrophage phenotypes or *E_e_* at either time point for either age group were used to define the ranges of each parameter during the next stage of sampling. Uniform sampling was used over this refined space for the second stage. The final step was iterative stochastic local search; using four iterations, successively restricting the space by adding more components of the criteria until in the end all remaining sets matched every criterion for one of the age groups.

In summary, the parameters generated by this sampling process underwent a selection process following this general outline for each parameter set: (1) determine the numerical steady state associated with the parameters in the absence of ventilation, (2) check whether the steady state variable values fell with in the range of the experimental data at time zero for either the old or the young experimental group, (3) if the resulting transient fell within the range for either group, simulate 2 hours of ventilation, and (4) check whether the resulting transients fell within the range of the experimental 2 hour data for the appropriate age group.

#### Classification, regression and sensitivity

Several methods were fit on various subsets of the parameter sets which fell in the range for either the old or young data. A suite of model fitting algorithms were applied using the R package H2O [79], including Generalized Linear Models, Distributed Random Forests, Gradient Boosting Machines, and Neural Networks. The best model in terms of 5-fold cross-validated F1 score (for classification) or root mean squared error (for regression) was chosen and the variable importance values were calculated for each.

Local sensitivity analysis was also used to measure the relative sensitivity of the model parameters for each experimental group. The methods were implemented using the SimBiology package available in MATLAB [77] which calculates the time-dependent derivatives of the model sensitivity to each parameter evaluated at specified time points. Details about the calculations performed can be found in Martins et. al [80]. Default settings were used for the sensitivity matrix. The overall relative sensitivities for each parameter were were calculated by taking the root mean square of the sensitivity values at the chosen time points for the chosen variables. The resulting values for each parameter are normalized by scaling each by the maximum overall sensitivity.

## Results/Discussion

### Experimental results

A large difference was observed in M0 cells at baseline between young and old mice (Fig. 2). There were increases in M1 marker expression in the alveolar macrophage populations (Fig. 2) from both the young and old mice after 2 hours of PCMV. M2 cells were increased at baseline in the old mice, with overlapping ranges in the PCMV mice. High PCMV enlarged the airspace in both young and old mice. The mean linear intercept, which is an index of airspace enlargement, was quantified to further assess the extent of injury (Fig. 2) . There was significantly increased airspace enlargement in the old PCMV group compared to the young and old non-ventilated controls. These findings suggest that there was a substantial generation of acute lung injury in both the young and old age groups; however, the severity appears to intensify with the old mice.

## In silico **results**

### Plausible parameter sets and transients

Iterative sampling used for the large sampling space produced 19,202 plausible parameter sets that fit both time points for either young or old experimental data. Of these sets, 17,477 fit the young data and 1,725 fit the old data. General behavior for the variables *E_h_*, *E_e_*, *M*_0_, *M*_1_, *M*_2_, *N*, and *AN* are shown in Fig. 3. Additionally, the percentage of the macrophage activity that was *M*_0_, *M*_1_, or *M*_2_ were plotted. The scaled version of each was compared to the experimental data.

**Fig 3.**
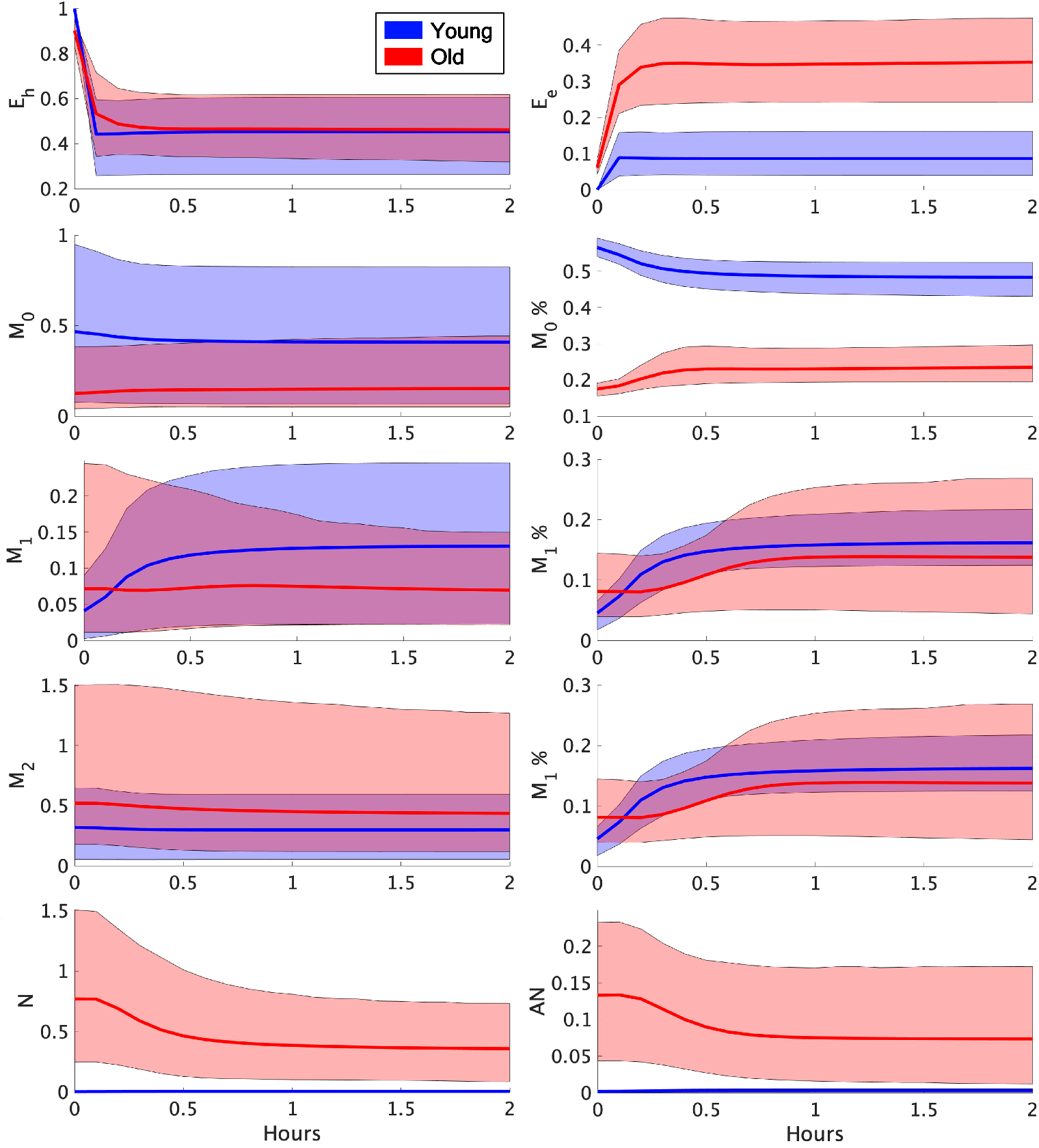
Old and young transients. The solid blue and red lines are plotted using the mean response for the transients associated with each experimental group at each time point. The surrounding bands mark the 10th and 90th percentile at each time point. M0%, M1%, or M2% represents the percentage of the macrophage activity that is M0, M1, and M2, respectively. These percentages along with the *E_e_* variable were compared to the experimental data.

Sets were classified using the proportion of their associated *E_h_*level before ventilation (*t* = 0) and directly after the 2 hour ventilation (*t* = 2). Sets were classified at each time point as healthy when *E_h_ >* 0.9, moderate when 0.5 *< E_h_ <* 0.9, and severe when *E_h_ <* 0.5. The resulting classifications are shown in Fig. 4.

**Fig 4.**
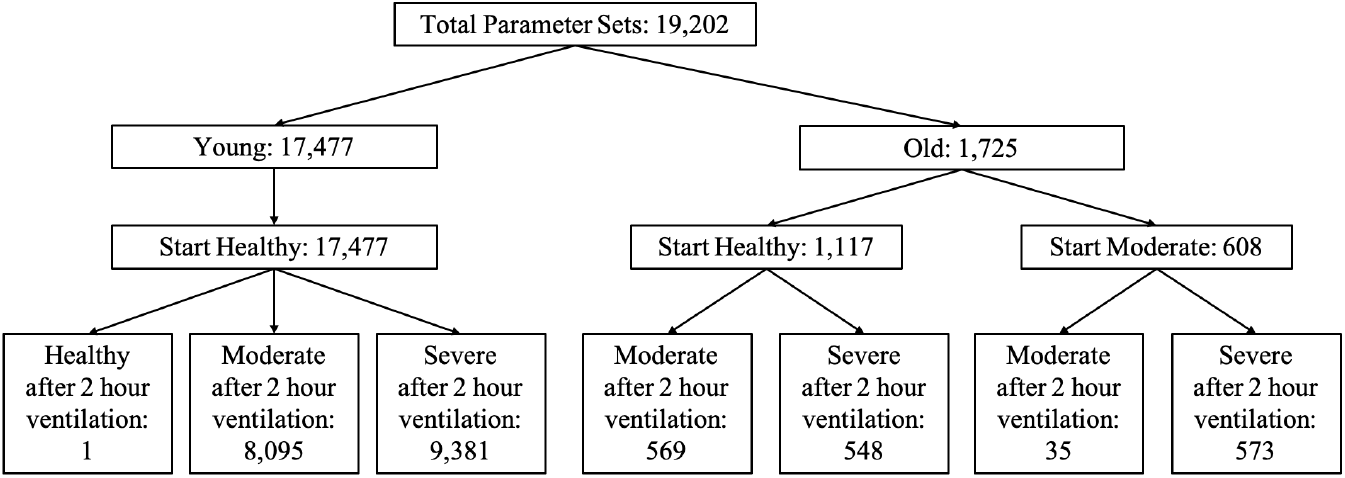
Classifications of plausible sets were made using the *E_h_* proportion before and after 2 hour ventilation. Transients were classified as healthy when *E_h_ >* 0.9, moderate when 0.5 *< E_h_ <* 0.9, and severe when *E_h_ <* 0.5.

This scheme created a natural division in the plausible parameter sets into the following groups by age and ventilation response: young sets that started healthy and became moderate after ventilation (Young H2M), young sets that started healthy and became severe after ventilation (Young H2S), old sets that started healthy and became moderate after ventilation (Old H2M), old sets that started healthy and became severe after ventilation (Old H2S), old sets that started moderate and stayed moderate after ventilation (Old M2M), and old sets that started moderate and became severe after ventilation (Old M2S). We will exclude the one young set that remained healthy after ventilation.

The experimental data was used as acceptance criteria, so these same differences are observable in the *in silico* model transients. This includes macrophage percentages and airspace enlargement. Neutrophil counts were not included in data collection but do exhibit an observable difference in the model simulations as seen in Fig. 3. The old transients exhibited much higher levels of neutrophils compared with the young sets. This is consistent with *in vivo* results where increased neutrophil infiltration and alveolar damage were correlated [81]. Higher neutrophil infiltration has also been observed in older individuals following lung injury [82].

#### Importance factors for age classification

Importance values for the various classification methods are shown in Fig. 5. Plot (a) exhibits ranked parameters in the classification of old and young parameter sets. In terms of scaled importance, the first four parameters are of interest since the numerical value decays significantly at the fifth ranked parameter. Thus, the following parameters had high importance in classifying between the old and young parameter sets: *k_em_*_1_, the rate of phagocytosis of damaged cells by *M*_1_, *x_er_*, a Hill-type constant regulating the effectiveness of repair of damaged cells by repair mediators, *k_m_*_1_, the rate of *M*_1_ migration into the alveolar compartment, which is novel to this iteration of the model, and *x_mne_*, a Hill-type constant regulating the effectiveness of macrophages and neutrophils to damage epithelial cells.

**Fig 5.**
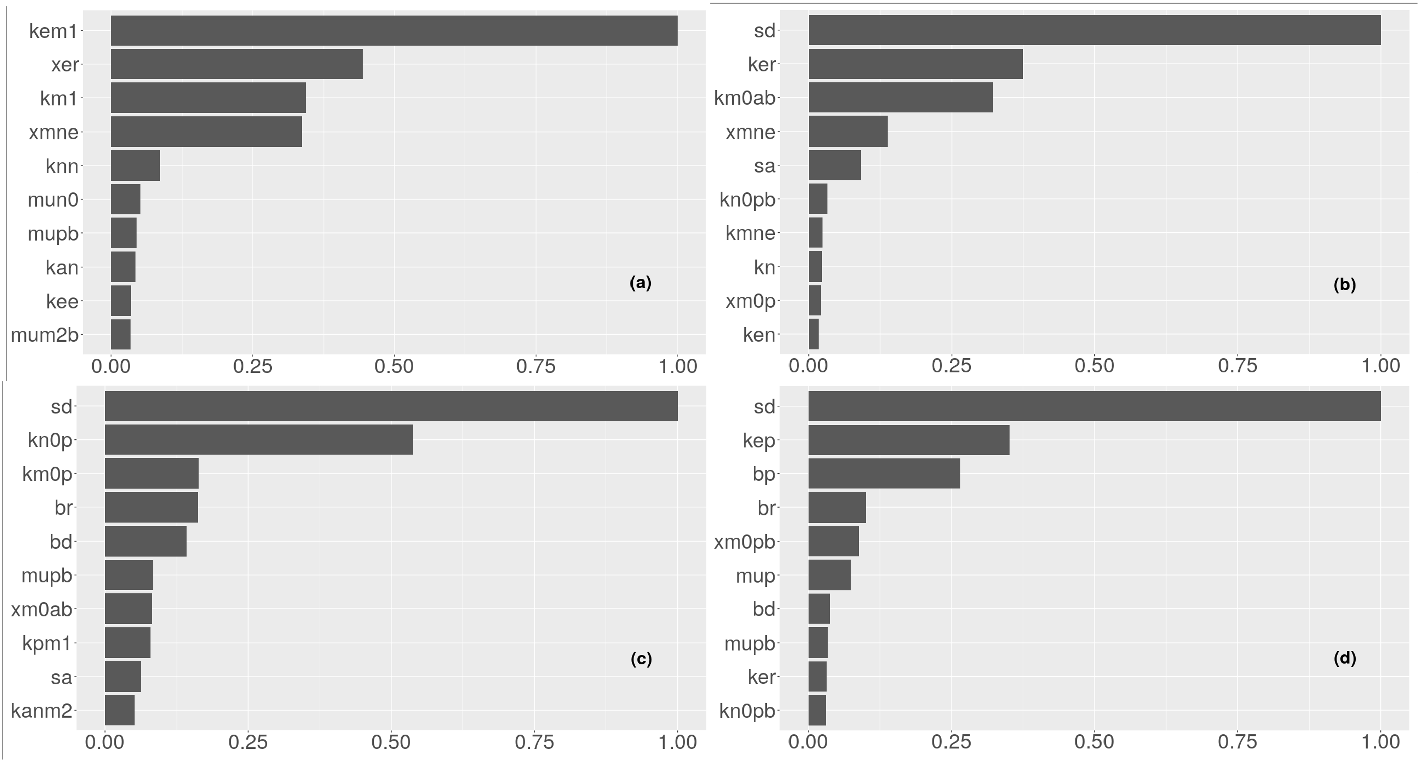
Scaled variable importance for top 10 predictors for (a) all observations (predicting young or old), (b) old class (predicting healthy or moderate at time 0), (c) healthy, young class (predicting moderate or severe after 2 hours), (d) healthy, old class (predicting moderate or severe after 2 hours)

The parameters *k_em_*_1_, *x_er_*, and *x_mne_*specifically involve repair and damage of the epithelial cells. The experimental data (Fig. 2, “Airspace Enlargment”) and *in silico* simulations (Fig. 3, “*E_e_*”) both exhibit a discrepancy in the epithelial variables. In terms of the actual data, the experimental range for airspace enlargement (Fig. 2, “Airspace Enlargement”, correlated with the model variable *E_e_*) does overlap between old and young subjects, but a general increase in this value is observed in the old mice. However, simulations show a clear distinction in the variable *E_e_* (Fig. 3, “*E_e_*”).

Therefore, the classification methods found parameters associated with epithelial dynamics important factors for determining the age group. Why other parameters affecting this variable were not found to be as important is unclear.

The parameters *k_m_*_1_, *k_em_*_1_, and *x_mne_*involve macrophages, specifically the M1 phenotype. This was not the expected phenotype to be associated with age classification given that the experimental groups have non-overlapping ranges for M0% and M2% at one or more time points (Fig. 2) and for the *in silico* simulations (Fig. 3). The *M* 1 phenotype changes are driven by the ventilation and their percentages are directly linked to the M0 and M2.

An additional classification suite was performed for the old parameter sets prior to ventilation, shown in Fig. 5, plot (b), since there was variety in the starting states of these simulations. The parameter *s_d_* holds the largest importance value as it contributes directly to damage to the epithelial cells. The parameters *k_er_*, the rate of repair to damaged cells by repair mediators, and *x_mne_*, a Hill-type constant regulating the effectiveness of macrophages and neutrophils to damage epithelial cells, are both direct parameters in the epithelial equations contributing to damage and repair processes. The parameters *k_m_*_0*ab*_, the rate of differentiation of *M*_0*b*_ by *a_b_*, and *s_a_*, the source rate of background *a_b_*, also had high importance factors. Both parameters are involved in the function of AIMs and have not yet been discussed in the topic of lung health classification. AIMs as well as their cellular counterparts, namely M2 macrophages, control the pro-inflammatory response and are crucial to preventing a feedback loop of chronic inflammation. As discussed earlier, both the pro- and anti-inflammatory responses are needed to ensure effective healing. Thus, despite their absence in the epithelial equations, AIMs are crucial to controlling cellular damage and promoting repair. The classification results reflect this relationship.

#### Importance factors for response

Plot (c) in Fig. 5 exhibits classification for a moderate or severe state after two hours of ventilation for young parameter sets. The most important parameters in classifying moderate or severe lung health were *s_d_*, the rate of epithelial damage from the ventilator, *k_n_*_0*p*_, the rate of activation of *N* in the alveolar compartment, novel to this iteration of the model, *k_m_*_0*p*_, the rate of differentiation of *M*_0_ by *p*, *b_r_*, baseline repair of damaged cells, and *b_d_*, baseline death rate for damaged cells.

For old sets that started healthy classification methods were used to determine importance factors for a moderate versus severe response to ventilation, Fig. 5(d). The most important parameters for classification were *s_d_*, *k_ep_*, the rate of self-resolving repair mediated by *p*, *b_p_*, baseline self-resolving repair of epithelial cells, *b_r_*, and *x_m_*_0*pb*_, a Hill-type constant regulating the effectiveness of differentiation of *M*_0*b*_ by *p_b_*.

The actual parameters with high importance factors differ between old and young parameter sets but generally involve rates of damage and repair. The parameter *s_d_* was the top parameter for each group. This parameter contributes directly to decreasing *E_h_* and thus influences the resulting classification of lung health. The parameters *b_r_*, *b_d_*, *k_ep_*, and *b_p_* all contribute directly to the epithelial variables and thus influence the level of *E_h_*as well. The remaining parameters *k_n_*_0*p*_, *k_m_*_0*p*_, and *x_m_*_0*pb*_ are not directly involved in the epithelial variables but do contribute indirectly to increased damage. All are involved in activation of inflammatory cells by PIMs. The pro-inflammatory phagocytic cells contribute to additional cellular damage. This is a well-established phenomenon of the inflammatory stage [83, 84].

#### Local sensitivity analysis for representative sets

Local sensitivity analysis was performed to measure the sensitivity of the variables *E_h_*and *E_e_* that are most important in determining the expected lung health of the simulations for the different groups. Sensitivity analysis was performed using user-defined representative sets.

A representative set was determined for each age and ventilation response group such that it was one of the parameters in the set and its behavior was most inline with the mean of the key variable transients for all the parameter sets in its group. The mean of the transients was calculated for each group using the average values at each time point for *E_h_*, *E_e_*, *M*_0_, *M*_1_, *M*_2_, and *N* . A representative set from each group was then chosen by minimizing the residual sum of squares for all six mean transients. The representative sets are plotted for each variable in Fig. 6. For each of the representative sets a local sensitivity analysis was performed and top ten sensitive parameters for each group are given in Fig. 7.

**Fig 6.**
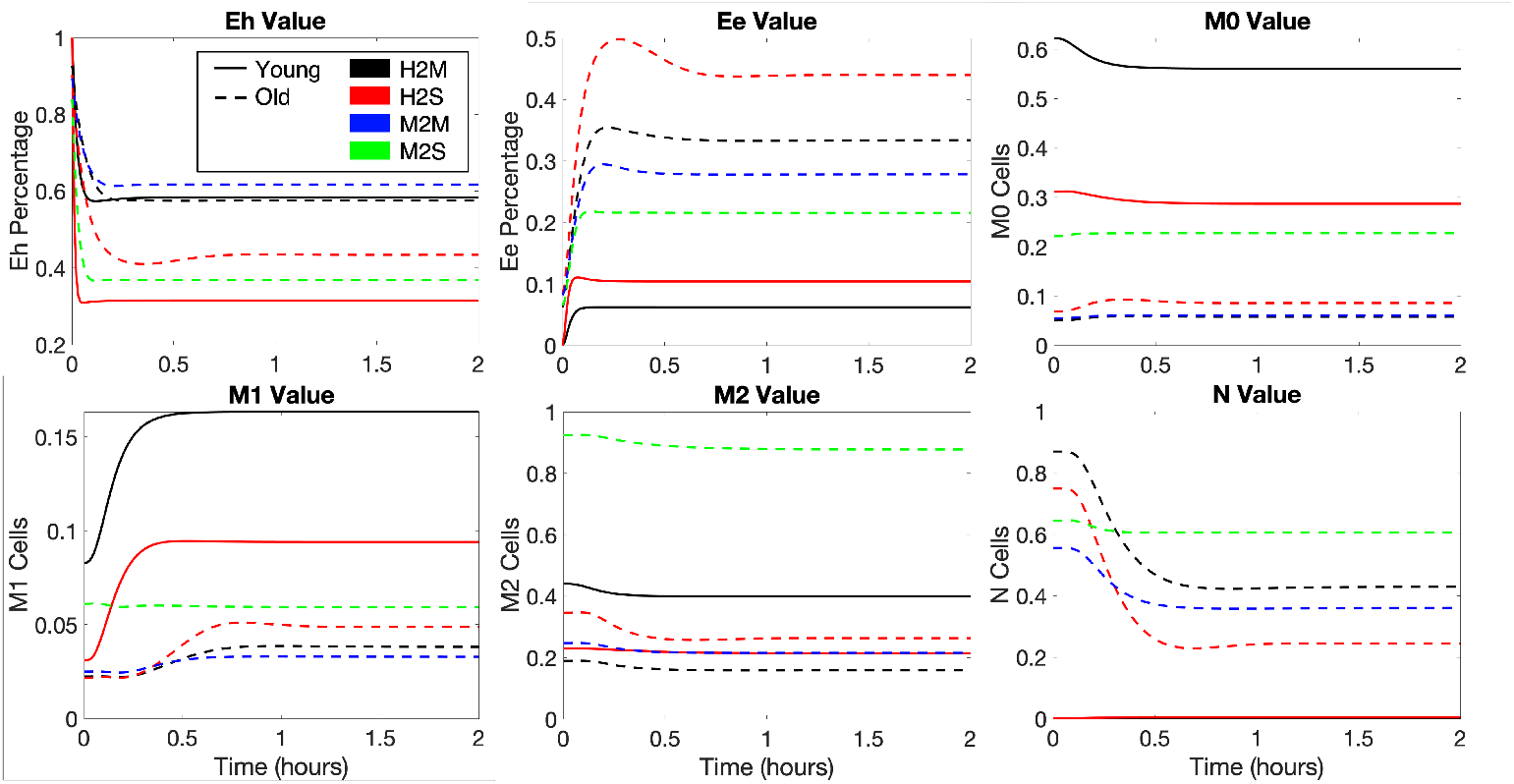
Representative sets for the various responses to ventilation using best fit sets to the mean response for each group. Transients are plotted for healthy epithelial cells (*E_h_*), dead epithelial cells/empty space (*E_e_*), naive macrophages (*M*_0_), M1 macrophages (*M*_1_), M2 macrophages (*M*_2_), and neutrophils (*N*).

**Fig 7.**
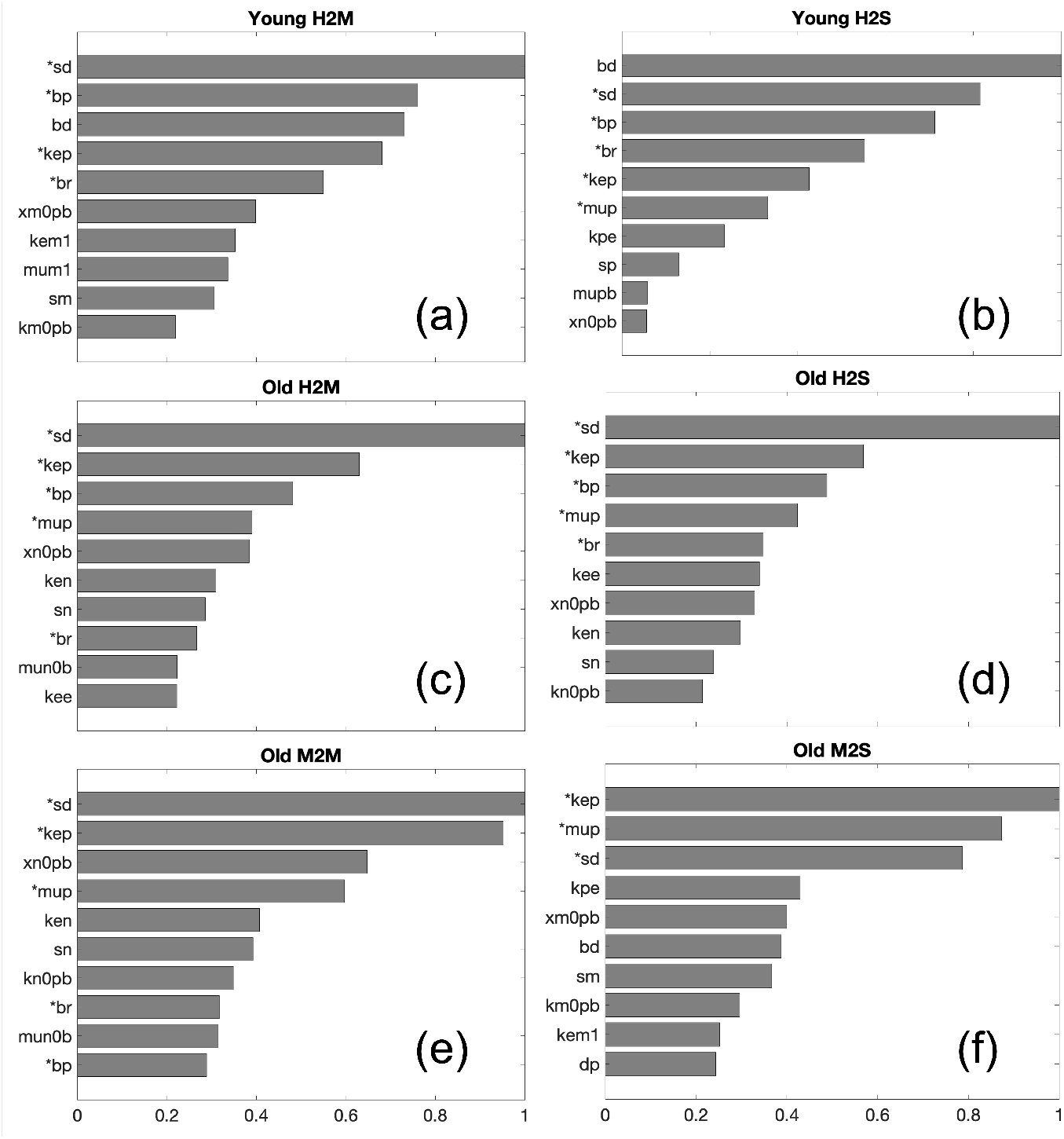
Normalized local sensitivity values for the top ten parameters in each representative set. Plots A-F are the sensitivities for the sets corresponding to Young H2M, Young H2S, Old H2M, Old H2S, Old M2M, and Old M2S, respectively. Parameters indicated with an asterisk had sensitivities greater than 10% of the maximum for all representative sets.

The model variables were considered to be sensitive to a parameter if the sensitivity value was larger than 10% of the maximum value. Multiple parameters were identified above this threshold for all of the parameter sets, namely; *b_p_*, baseline self-resolving repair of epithelial cells, *s_d_*, the rate of damage from the ventilator, *b_r_*, baseline repair of damaged cells, *k_ep_*, the rate of self-resolving repair mediated by *p*, and *µ_p_*, decay rate of *p*. These parameters mainly affect accumulation of damage, ability to repair, and dynamics involving PIMs. Since *E_h_* and *E_e_* were the variables used in the sensitivity calculations, it is unsurprising that the parameters in those equations had high sensitivity values. We also again see parameters involved in the pro-inflammatory response to be important.

Additional parameters were found to be sensitive in the majority of the groups. Parameters *s_p_*, source of PIMs, *s_n_*, source of neutrophils, and *k_en_*, rate of phagocytosis of damaged cells by neutrophils, had sensitivity values larger than 10% of the maximum in all groups expect Young H2M. The parameter *x_n_*_0*pb*_, a Hill-type parameters involved in the activation of neutrophils by PIM, had a sensitivity value larger than 10% of the maximum in all the old groups. Again, we see parameters involved in PIMs and cell production as well as their processes. Interestingly, the parameter *k_em_*_1_, phagocytosis of damaged cells by M1 macrophages, had small sensitivity values for the majority of the groups. Since we are only looking at time points contained within the 2 hour ventilation time, this could explain why parameters involving macrophages were considered insensitive. Neutrophils are typically the first cells to arrive, while macrophages generally arrive later in the inflammatory stage. Additionally, previous research has hypothesized that a sustained inflammatory insult is primarily mediated by neutrophils, and is particularly important in the development of acute lung injury and ARDS [85].

#### Modulating response to ventilation

The results of the local sensitivity analysis were used to simulate a pseudo-intervention for the parameter sets associated with a severe state after 2 hours of ventilation: Young H2S, Old H2S, and Old M2S.

The parameters chosen as influential across all groups from the sensitivity analysis were *b_p_*, *s_d_*, *b_r_*, *k_ep_*, and *µ_p_*. These parameters were increased by 10% one at a time and the percent increase or decrease in the variables *E_h_*and *E_e_* at 2 hours was calculated for each parameters set within the Young H2S, Old H2S, and Old M2S groups. Table 3 shows the minimum, mean, and maximum percent change for each variable for each of these three groups. Supplementary materials includes this same analysis with the additional parameters identified as influential in the majority of the groups as well as an intervention starting after one hour of ventilation rather than prior to ventilation.

**Table 3.**
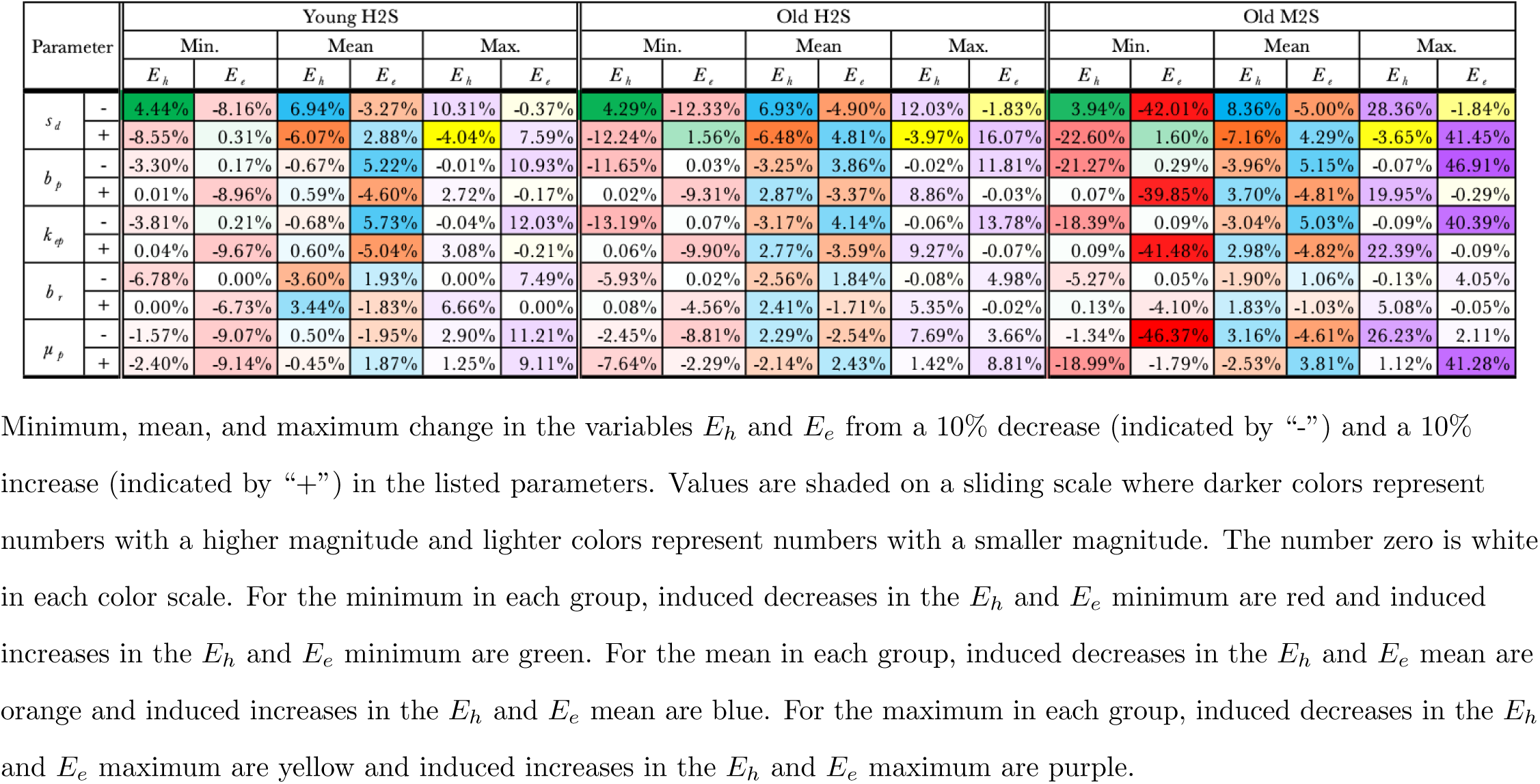
Variable effects of modulating parameters.

The intent of modulating parameters is to explore potential targets for therapeutic interventions prior to or during MV. The results in Table 3 demonstrate the various outcomes as well as differences among groups. A range of responses is expected in practice since patients have unique health profiles. However, the mean provides a reasonable expectation for the average response. The results shown in Table 3 exhibit the wide range of behavior possible with the model and parameter sets and specific groups and parameter combinations produced large ranges of possible values.

Specifically, the Old M2S parameter sets had a high level of variability in the observed maximums, especially for the variable *E_e_*. The parameters *b_p_*, *k_ep_*, *s_d_*, and *µ_p_* produced a possible difference of around 40-50% in the variable *E_e_*. Potential large decreases were also observed for the same parameters producing a decrease in *E_e_* at similar magnitudes. Generally, overall averages were not as considerable and were all less than 10% in absolute value, with some producing average differences close to zero.

Parameters had varying outcomes on the variables, with some increasing *E_h_* while simultaneously increasing *E_e_*; thus not necessarily improving health outcome. However, an increase in the parameters *b_p_*, *b_r_*, and *k_ep_*consistently increased *E_h_* and decreased *E_e_*with a decrease in the parameter producing the opposite effect. The parameter *s_d_* exhibited the inverse relationship where an increase generally produced a decrease in *E_h_*and increase in *E_e_* with a decrease producing the opposite effect. It is clear by the color intensities that variation in the parameters generally had the most impact on the parameter sets in the Old M2S group. The largest impact was specifically observed in the variable *E_e_*. Since *E_h_* was used to define health states, a large change in *E_e_* may not result in a change in the health state as we have defined it.

## Conclusion

Age-dependent responses to ventilation are of medical interest given the increased need for ventilation and mortality rates of ventilated patients with age. Using mathematical modeling and statistical methods, we analyzed plausible ventilator responses associated with experimental groups for young and old mice with 2 hours of ventilation using macrophage phenotype and lung integrity data.

Classification results revealed that parameters involved in repair and damage of epithelial cells as well as parameters involved in macrophage function were important in separating the old and young parameter sets. The parameters involved in repair and damage of epithelial cells were expected results given the discrepancies observed in the airspace enlargement variable of the experimental data as well as past research that has shown poorer lung health in aged subjects. Parameters involved in macrophage function were also expected to be associated with differences in old and young parameter sets based on past research; however classification results highlighted macrophage parameters specifically relating to the M1 phenotype which was not observed to have discernible differences in the experimental data. Disparities in the experimental data were instead observed in M0 and M2 macrophages at zero hours.

Classification results based on lung health state identified parameters involved in activation of inflammatory cells and mediators, and parameters involved in damage and repair to be important. These results are expected since repair and damage directly contribute to the variables used to classify the health state and processes involved in the pro-inflammatory response are known to affect local tissue health. Local sensitivity results identified similar parameters involved in damage and repair as we well as parameters directly related to a pro-inflammatory response, namely PIMs and neutrophils. Past research has identified a pro-inflammatory response, mainly mediated by neutrophils, to be significant in the development of acute lung injury and ARDS.

To explore how targeted interventions could change the poor responses to ventilation we modulated parameters that model outputs were sensitive to and evaluated changes in the model-predicted epithelial cell health. The local sensitivity results were used to select parameters to modulate and simulations were performed for all parameter sets in each of the representative groups with a severe health state after ventilation. In some cases, a wide range of responses was observed. The greatest effect was observed in the old representative set, specifically the variable *E_e_*, that started moderate and was severe after two hours. Some parameter modulations produced a potential increase or decrease in *E_e_* of about 40-50%. Overall, the mean response to modulation of each parameter had a magnitude less than 10%, with some near 0. Despite this, the targeted parameters offer potentially large improvements, particularly in the old parameter sets with poorer lung health. Additionally, and intervention performed mid-way through ventilation also produced potential improvements in the transients but to a lesser extent.

The exploration of age differences in VILI expresses early attempts to create more personalized medical interventions. Despite the observed differences in MV response with age, clinical use and interventions for MV remain a one-size-fits-all approach. Understanding the underlying differences in the immunological response to VILI could allow for better, targeted interventions. This model can also be adapted to account for non-VILI associated lung injury, such as an infection or inhaled toxins. The statistical and mathematical method used with this model can then determine key components of the immune and repair responses for those insults. Insults could be combined with ventilation to better explore the age-dependent response to VILI while accounting for co-factors that lead to the necessity for ventilation.

## Supplementary Materials

### S1 Supplementary table

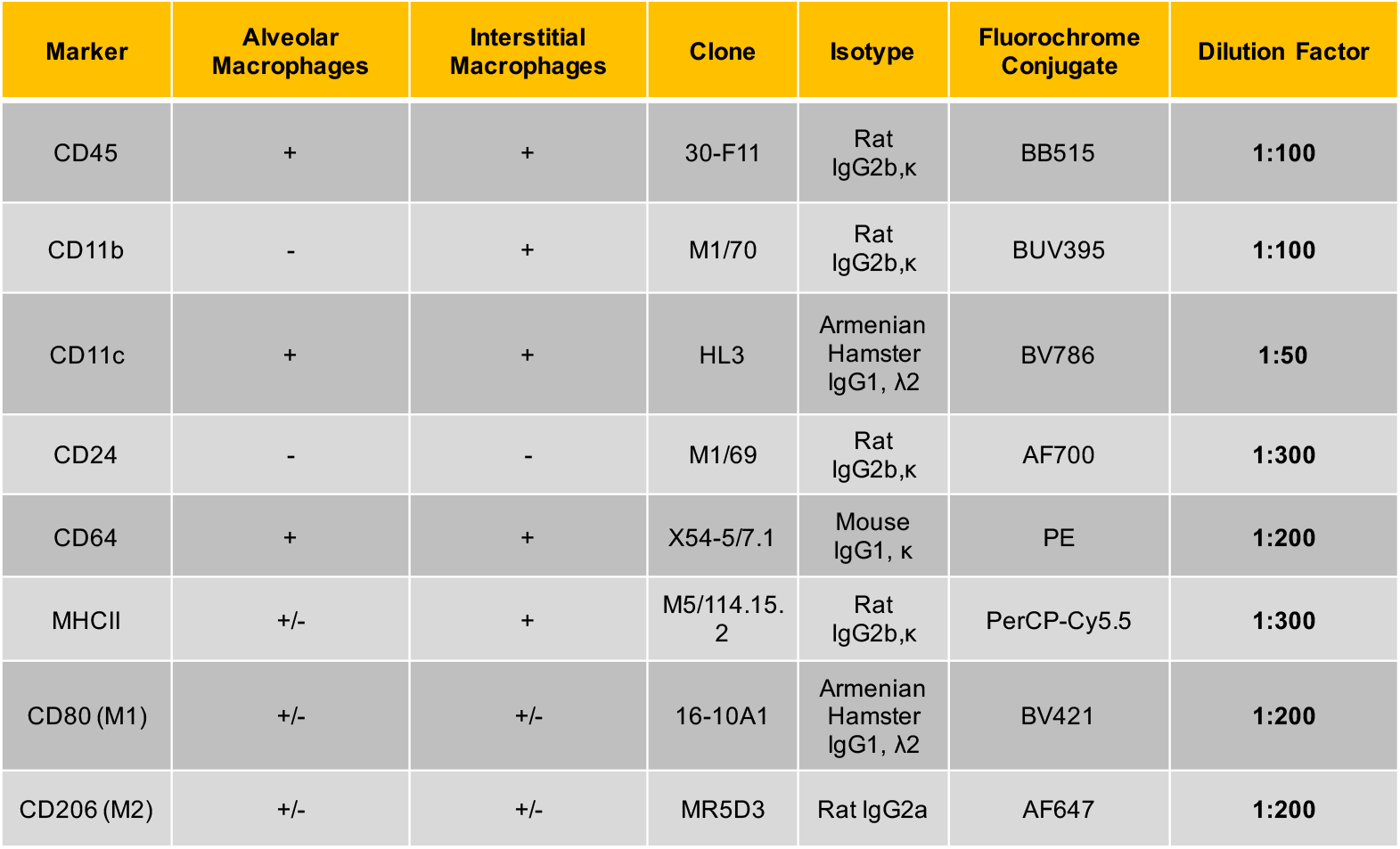

### S2 M0 macrophage equations

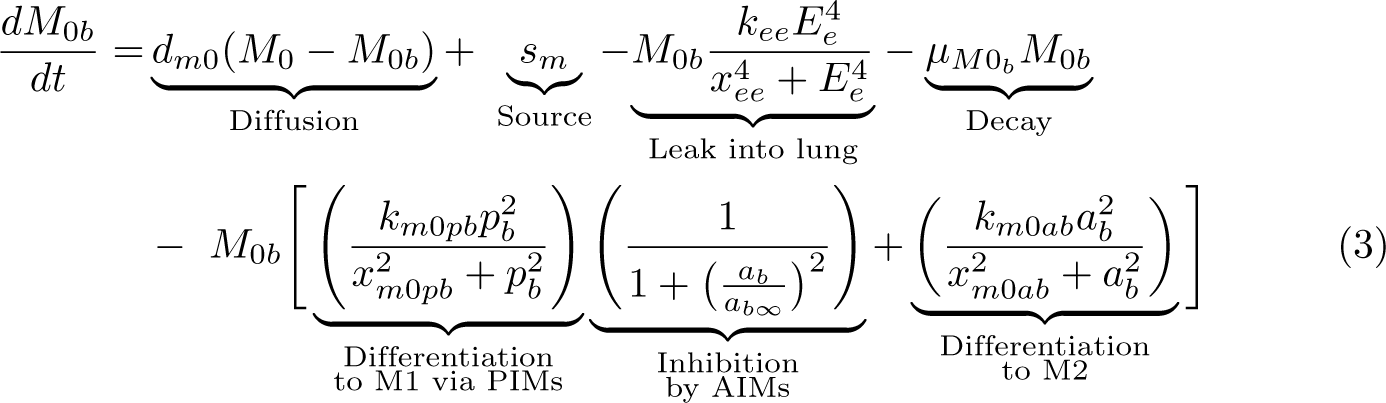

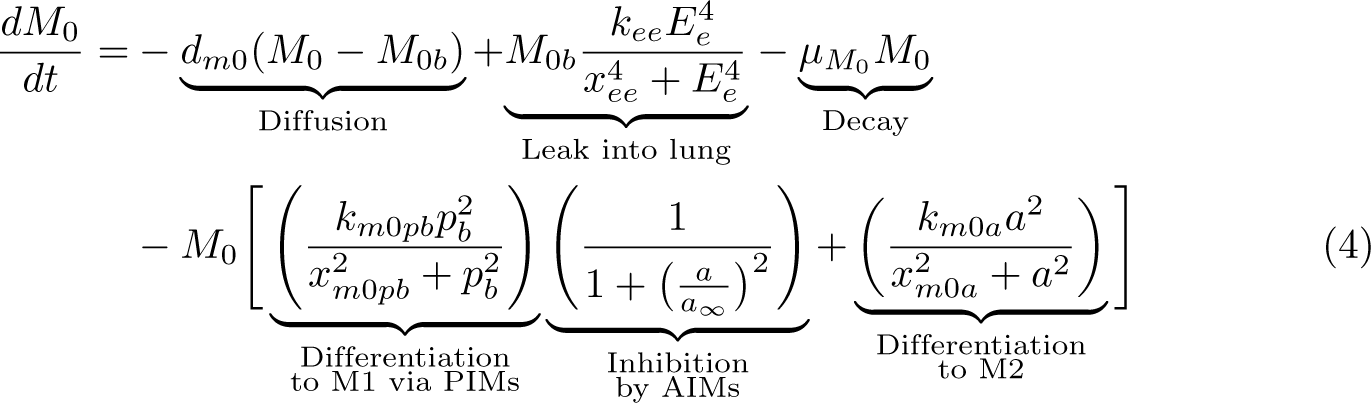

### S3 M1 macrophage equations

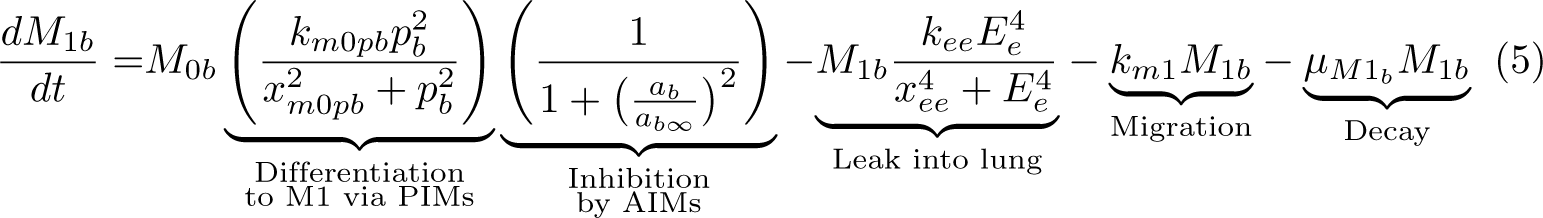

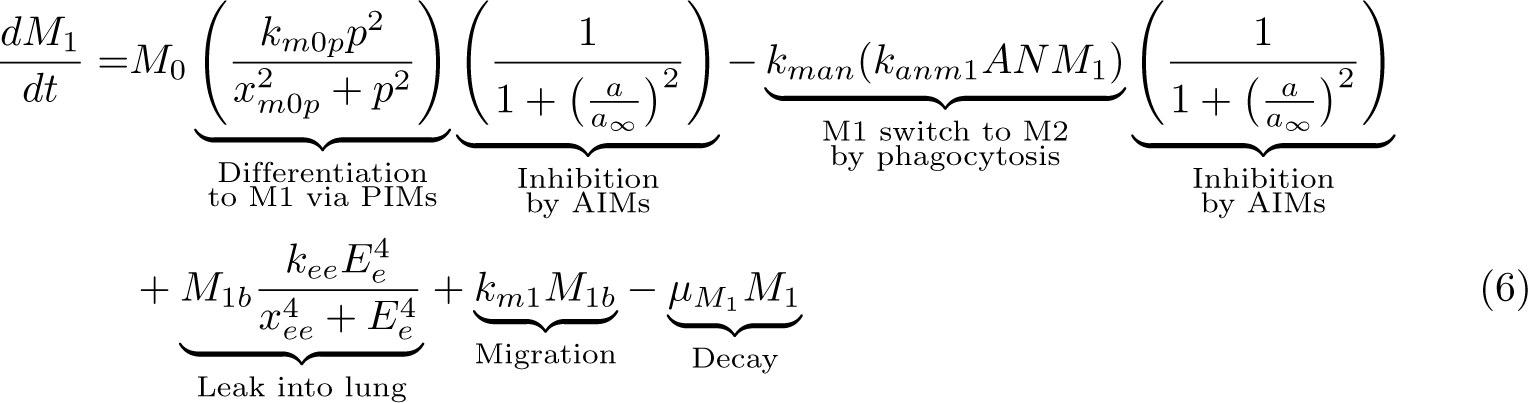

### S4 M2 macrophage equations

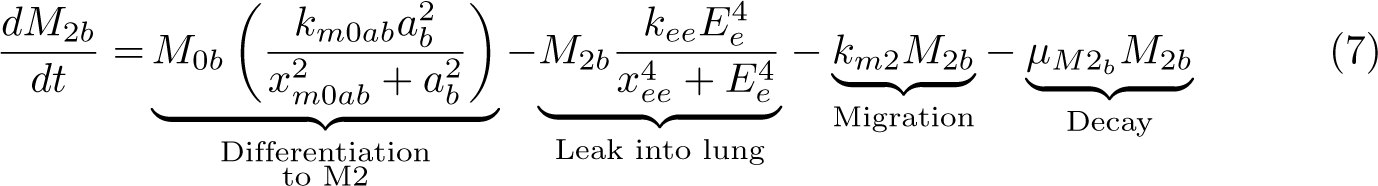

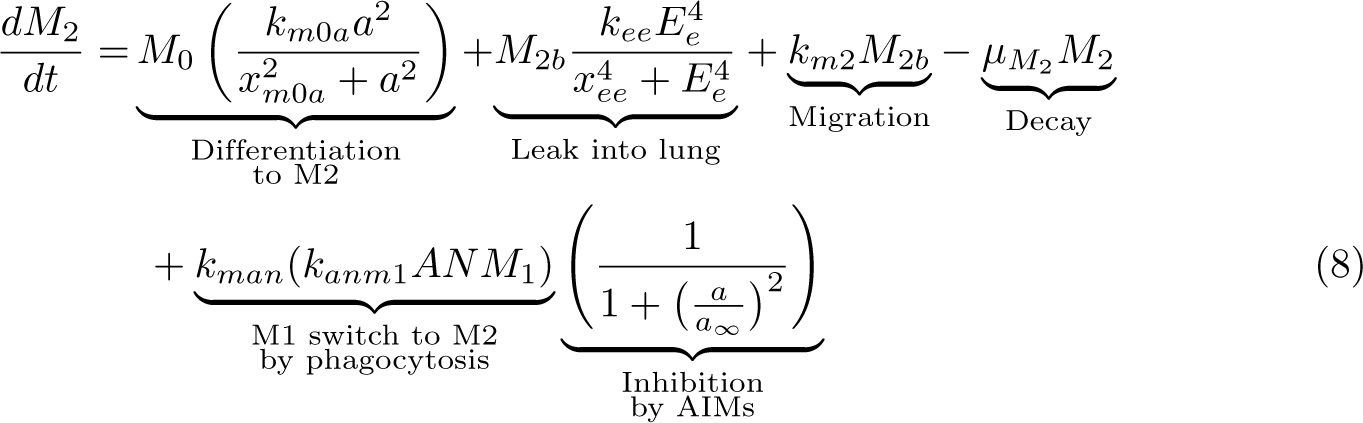

### S5 Neutrophil equations

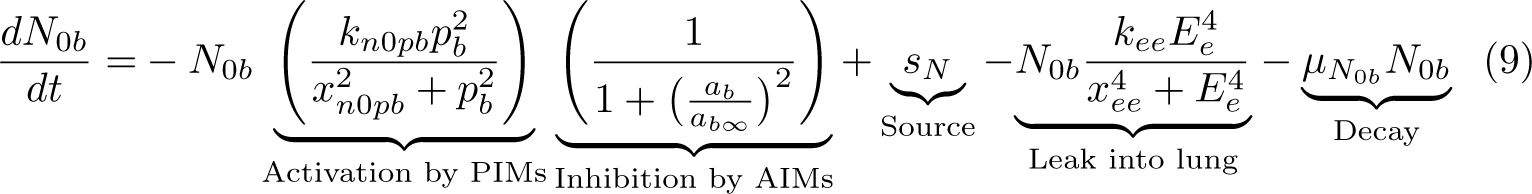

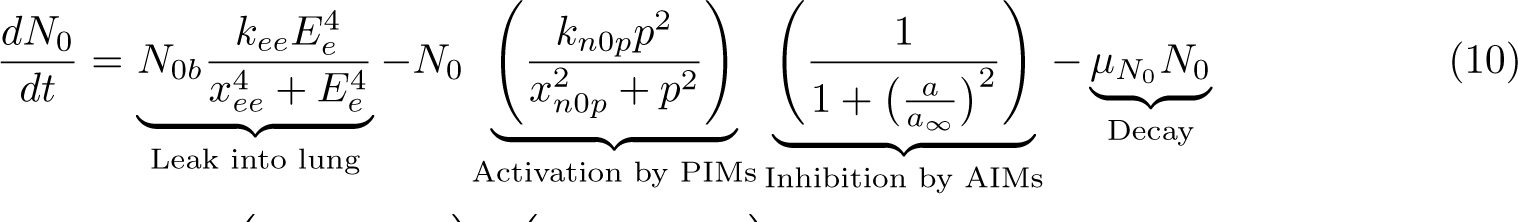

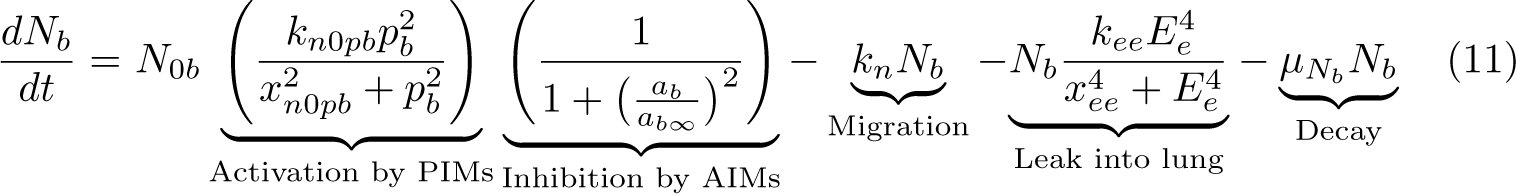

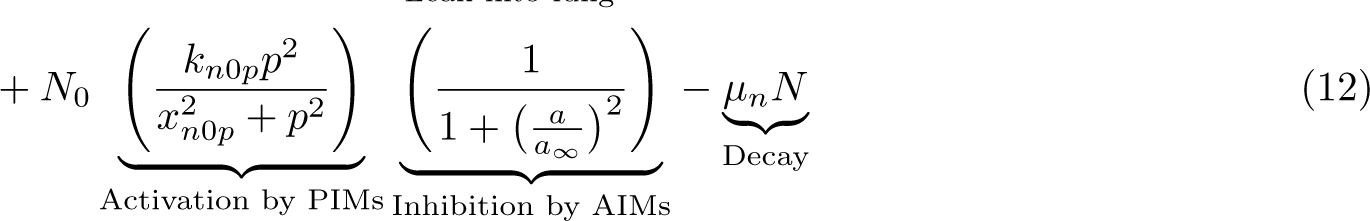

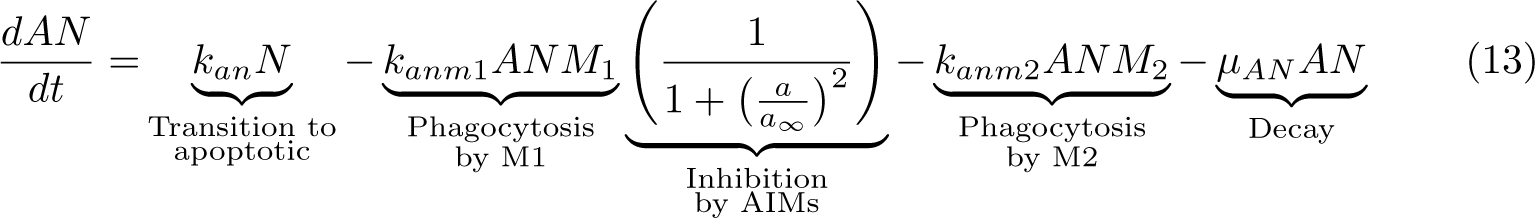

### S6 Pro- and anti-inflammatory mediators equations

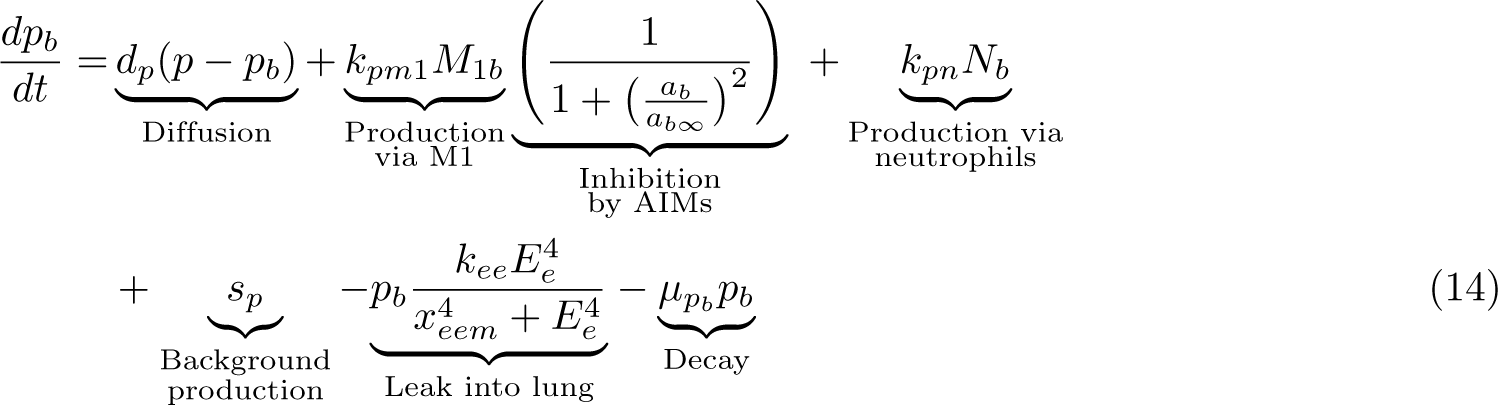

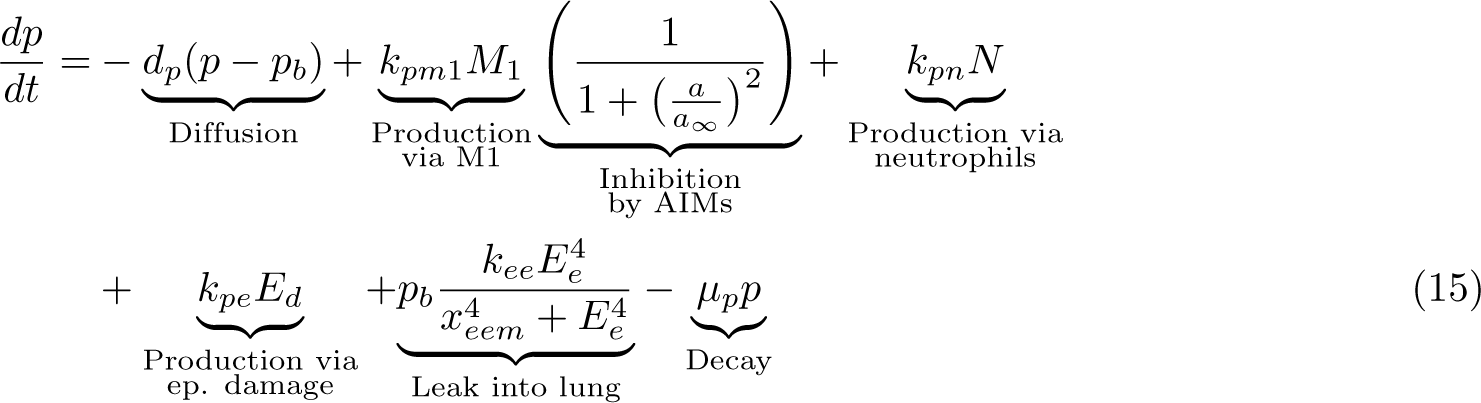

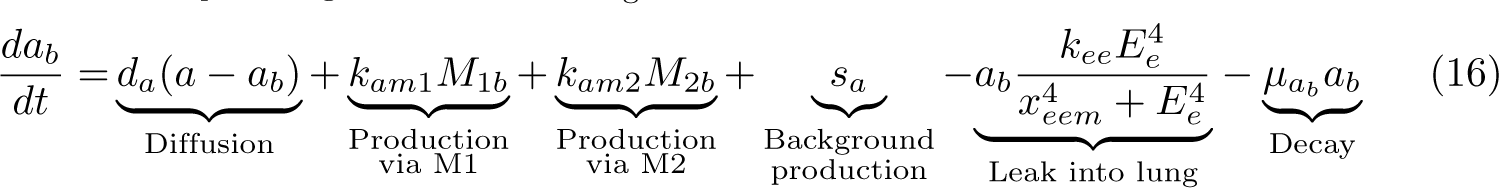

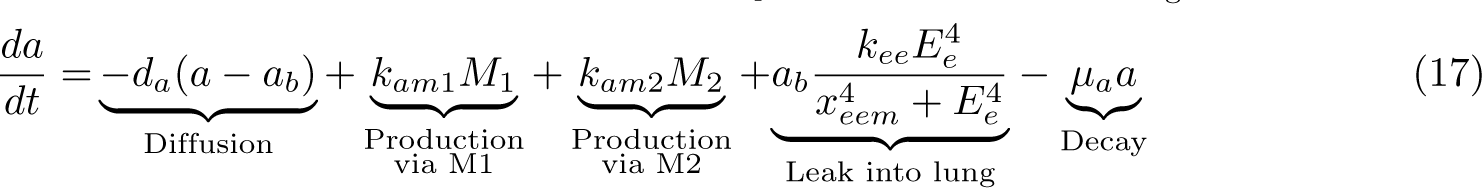

### S7 Repair and epithelial equations

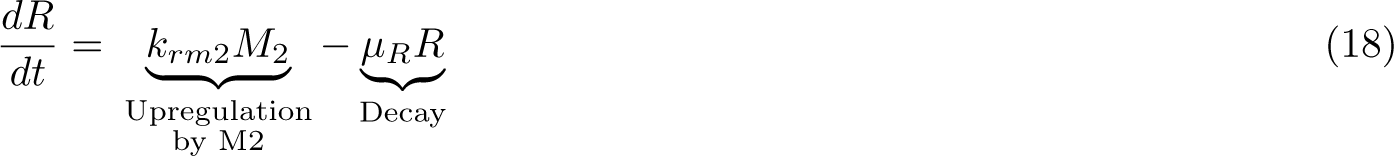

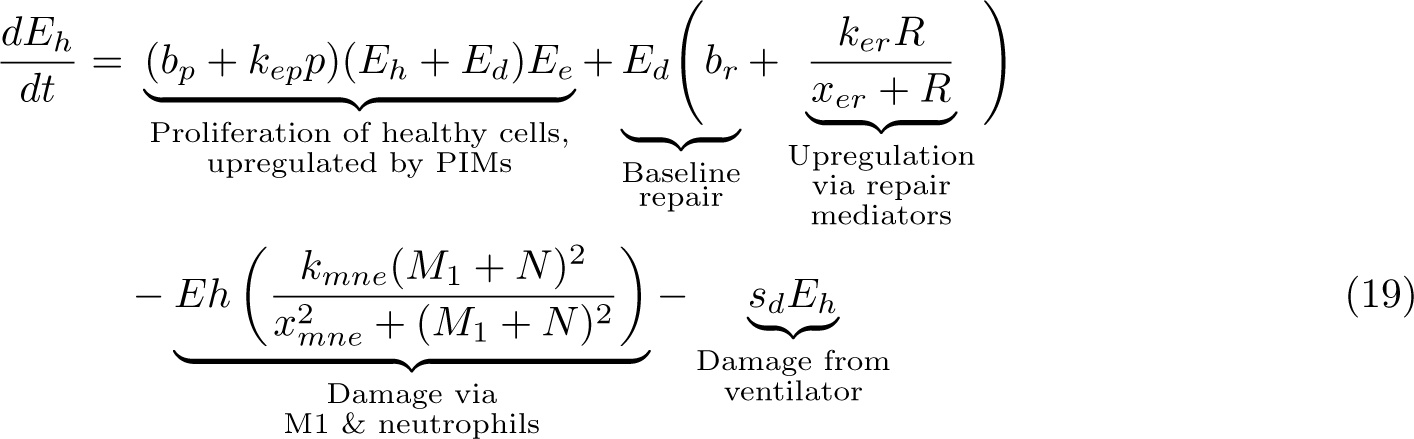

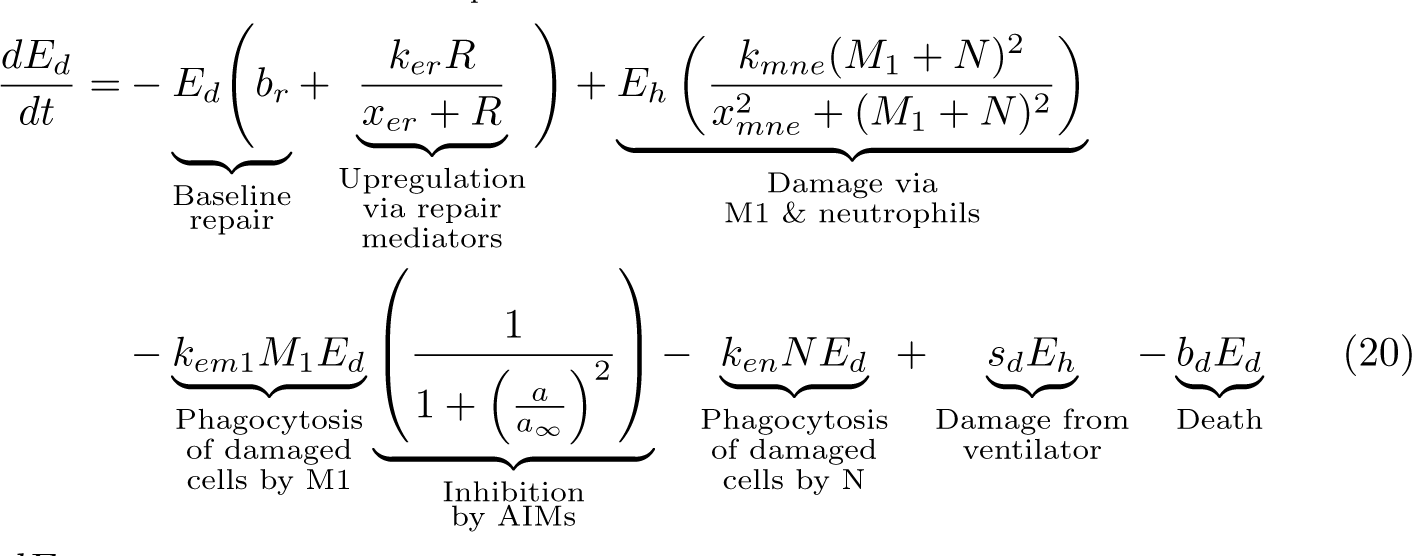

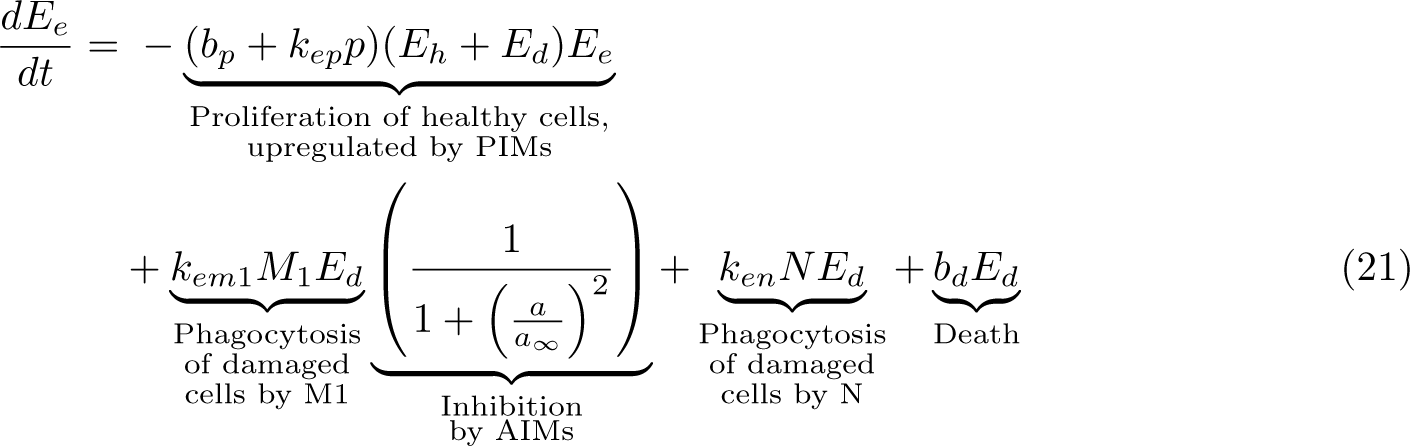

### S8 Full table modulating parameters prior to ventilation

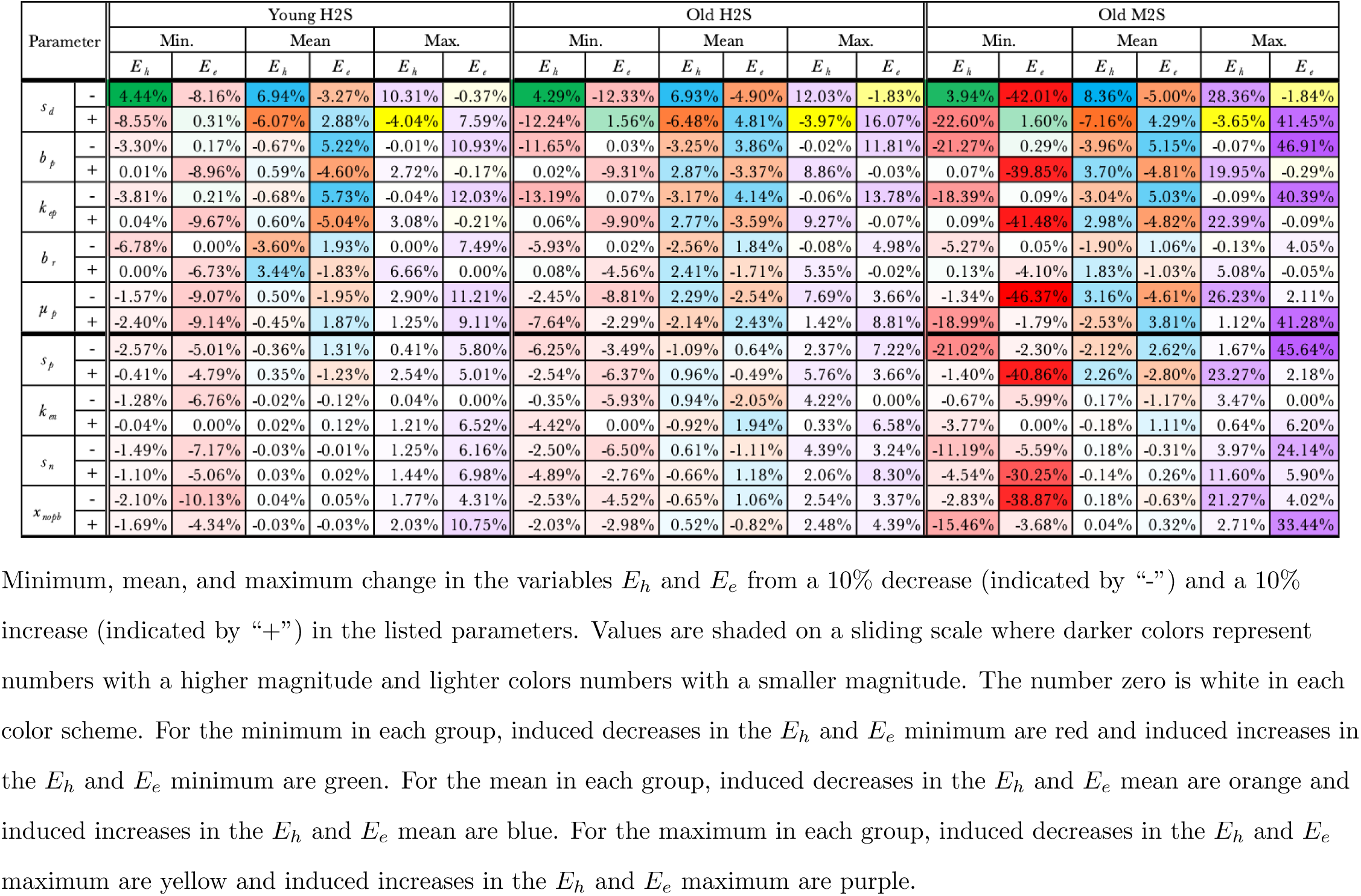

### S9 Full table modulating parameters after 1 hour of ventilation

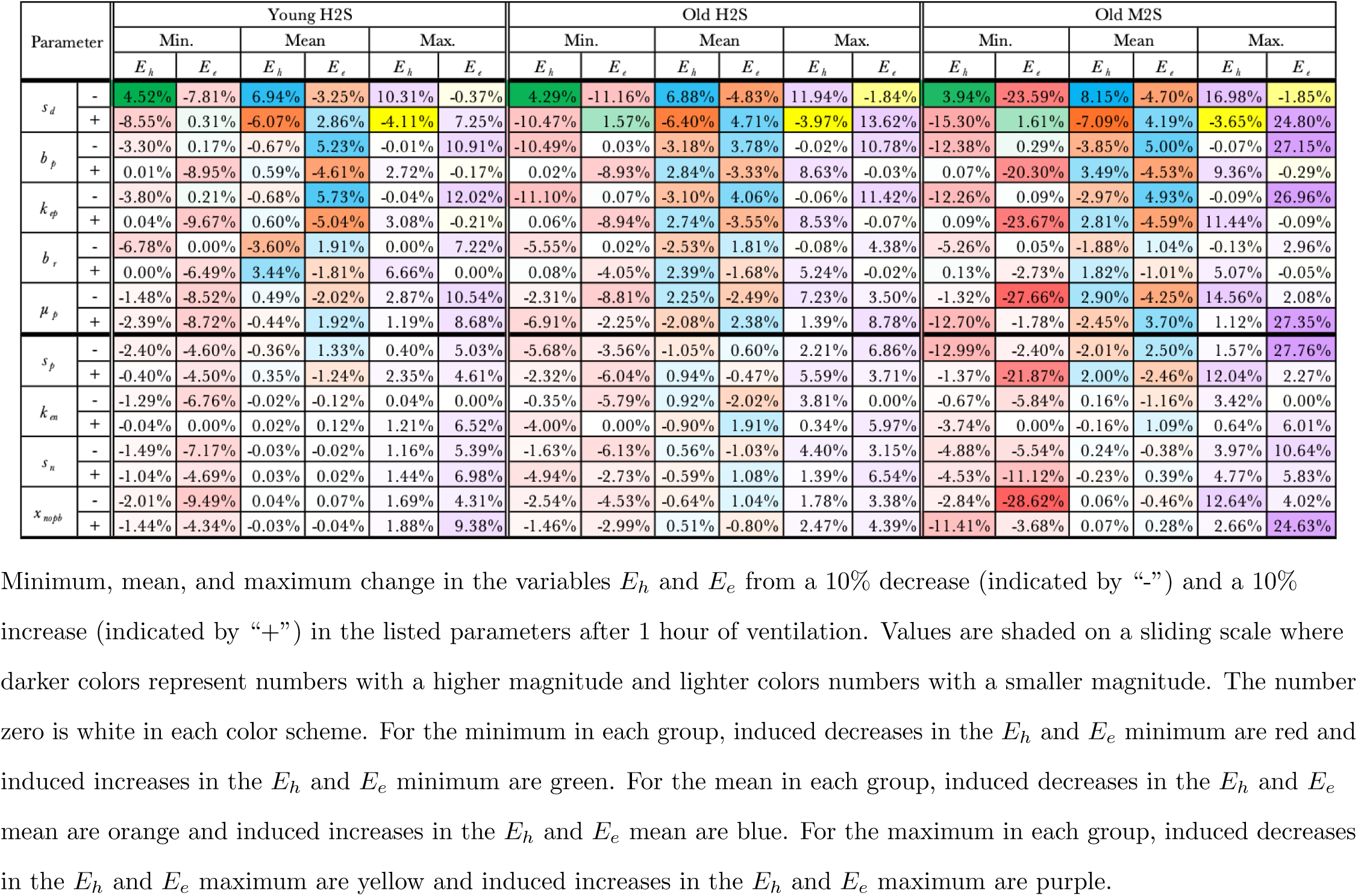

